# Dual role of brain endothelial Gpr126 in blood-brain barrier development and ischemic stroke

**DOI:** 10.1101/2022.09.09.507316

**Authors:** Nikolaos Kakogiannos, Anna Agata Scalise, Emanuele Martini, Claudio Maderna, Serena Magni, Giorgia Serena Gullotta, Maria Grazia Lampugnani, Fabio Iannelli, Galina V. Beznoussenko, Alexander A. Mironov, Camilla Cerutti, Katie Bentley, Andrew Philippides, Federica Zanardi, Marco Bacigaluppi, Gianvito Martino, Elisabetta Dejana, Monica Giannotta

**Author notes:** Correspondence to: Monica Giannotta.

## Abstract

The blood–brain barrier (BBB) acquires unique properties for regulation of the neuronal function during development. The genesis of the BBB coupled with angiogenesis is orchestrated by the Wnt/β-catenin signaling pathway. Aside from the importance of Wnt/β-catenin signaling, the molecular mechanisms that regulate these processes are poorly understood. Here, we identify the brain endothelial adhesion G-protein–coupled receptor Gpr126 as a novel target gene of Wnt/β-catenin signaling that is required for postnatal BBB development, and its expression is detrimental for ischemic stroke in adults. We show that Gpr126 expression is high in mouse brain endothelium during BBB formation, but decreases in the adult. Inactivation of Gpr126 in postnatal endothelial cells results in vessel enlargement and impairs acquisition of the BBB characteristics, such as increased neurovascular permeability, and reduced basement membrane protein deposition and pericyte coverage. Mechanistically, Gpr126 is required during developmental angiogenesis to promote endothelial cell migration, acting via an interaction between Lrp1 and α3β1-integrin, which couples vessel morphogenesis to BBB formation. Interestingly, in adult mice with an established BBB, the lack of Gpr126 expression in acute ischemic stroke is protective and coupled with reduced microglia activation, which contributes to an improved neurological outcome. These data identify Gpr126 as a promising therapeutic target to treat ischemic stroke.

## Introduction

The blood-brain barrier (BBB) separates the neural tissue from the blood circulation. The BBB is formed by a single layer of the endothelial cells (ECs) that line the blood vessel wall, which are surrounded by pericytes, astrocytes and mural cells that are embedded in the basement membrane (BM). Together, these form a structure that is commonly known as the ‘neurovascular unit’ (NVU) (Abbott et al., 2006; Zlokovic, 2011). The BBB ECs and all the cells that form the NVU maintain the homeostatic milieu to allow correct neuronal function and synaptic transmission.

Impairment of the BBB and its increased permeability can lead to pathological conditions, such as stroke, multiple sclerosis, HIV encephalitis, age-related dementia, and Alzheimer’s disease (Abbott et al., 2006). The BBB ECs form an active permeability barrier with unique biological features and transport system, as compared to ECs that form the vasculature in other organs. First, BBB ECs express specialized tight junctions (TJs) that strictly regulate the passage of molecules from the bloodstream into the interstitial brain tissue. TJs include the claudins, occludin, and junctional adhesion molecules that are connected to the cellular actin cytoskeleton via zonula occludens proteins such as ZO-1 (Berndt et al., 2019; Greene and Campbell, 2016; Irudayanathan et al., 2016). Then, adherens junction proteins, such as VE-cadherin, further strengthen the bonds between neighboring ECs (Brown and Davis, 2002; Stamatovic et al., 2016). Secondly, BBB ECs express exclusive transporters to regulate the dynamic influx and efflux of specific substrates (Zhao et al., 2015). For example, low-density lipoprotein receptor-related protein-1 (Lrp1) is a transporter of larger peptides and proteins, and it mediates the crossing of amyloid-β (a peptide secreted during Alzheimer’s disease) from the brain parenchyma into the circulation (Storck et al., 2016). Thirdly, BBB ECs have very low rates of transcellular vesicle trafficking, termed transcytosis, which limits transcellular transport through the vessel wall. Finally, to inhibit immune cell entry into the brain, BBB ECs express low levels of leukocyte adhesion molecules (Ransohoff and Engelhardt, 2012).

However, these specific barrier properties are not constitutive to ECs. Indeed, when brain ECs first enter the central nervous system (CNS) they do not inherently display barrier properties. Therefore, BBB formation is a gradual process that starts during embryonic development, with initial brain angiogenesis at embryonic stage-day E 9.5, and it is completed during early postnatal stages, up to 25 days in mice (Biswas et al., 2020). The BBB properties are also acquired by close interactions and cross-talk between ECs and the surrounding cells that form the NVU (Abbott et al., 2006; Zlokovic, 2011).

The NVU BM provides structural support for the cells, and it is a hub for intercellular communication and signaling. It is composed of structural proteins, such as collagen IV, fibronectin, laminins, and other glycoproteins. Collagen IV and fibronectin are secreted by ECs, pericytes, and astrocytes, and the depletion of either of these is embryonic lethal (Poschl et al., 2004; Theocharis et al., 2016). Laminins have several isoforms, and their appropriate balance is important in vessel and BBB formation (Menezes et al., 2014; Thyboll et al., 2002).

The BBB is neither absolute nor static, but rather dynamic and highly regulated by interactions between its cellular and BM components, along with their receptors, such as the integrins (Carvey et al., 2009; Neuwelt et al., 2011). Different β1-integrin isoforms are expressed by brain capillary ECs (α1β1, α3β1, α6β1, αvβ1) (Engelhardt, 2011; Milner and Campbell, 2002, 2006; Paulus and Jellinger, 1993), pericytes (α4β1) (Grazioli et al., 2006), and astrocytes (α1β1, α5β1, α6β1) (Milner R and Campbell 2006), and these bind to the BM and trigger signaling cascades to regulate cell survival, proliferation and migration.

Canonical Wnt signaling is the major pathway that regulates the onset of brain angiogenesis and BBB formation. Wnt ligands secreted by neural cells elicit canonical Wnt signaling in the endothelium by binding to the frizzled receptors (Wang et al., 2012) and the co-receptors LRP5/6, which in turn activate β-catenin–dependent pathways (Daneman et al., 2009; Liebner et al., 2008; Zhou et al., 2014). Activation of Wnt/β-catenin signaling influences TJs and BBB formation by increasing the expression of barrier-related genes, such as solute carrier family 2 member 1 (*Slc2a1*) and claudin-5 (*Cldn5*) (Daneman et al., 2009; Liebner et al., 2008). Deficiency of the endothelial Wnt/β-catenin signaling pathway affects cerebrovascular and BBB development, but does not affect the formation and function of other organs and tissues (Cullen et al., 2011; Daneman et al., 2009; Stenman et al., 2008).

Therefore, we hypothesized that the molecular targets of Wnt/β-catenin signaling in brain ECs might represent novel candidates for the control of BBB development. We and others have shown that several factors downstream of Wnt/β-catenin signaling promote BBB development, such as SRY-box transcription factor 17 (Sox17) (Corada et al., 2019; Corada et al., 2013) and fibroblast growth factor binding protein 1 (Fgfbp1) (Cottarelli et al., 2020). Further, forkhead box F2 (Foxf2) (Hupe et al., 2017; Sabbagh et al., 2018) maintains the BBB characteristics. However, the mechanisms of how canonical Wnt activation controls the formation of a functional BBB and how it links brain angiogenesis to BBB genesis are poorly understood. To address these, we activated Wnt/β-catenin signaling in murine ECs isolated from the brain microvasculature under treatment with the morphogen Wnt3a (wingless-type MMTV integration site family, member 3a). Microarray analyses unveiled the adhesion G-protein–coupled receptor (GPCR) 126 (*Gpr126*) as a novel gene regulated by Wnt/β-catenin signaling. Gpr126 is a member of the adhesion subfamily of GPCRs and it shares a seven-transmembrane domain with other GPCRs. However, Gpr126 is distinct in terms of its long, heavily glycosylated N-terminal regions that contain different adhesion domains that are separated from the seven-transmembrane-domain helix by a GPCR autoproteolysis-inducing domain (GAIN). Gpr126 can be cleaved at a GPCR proteolytic site in the GAIN domain to produce the N-terminal fragment and the C-terminal fragment (Patra et al., 2014). While the N-terminal fragment of GPR126 is known to form docking sites for extracellular matrix proteins, the C-terminal fragment of GPR126 can engage α subunits of heterotrimeric G-proteins to transduce extracellular stimuli into the intracellular milieu (Liebscher et al., 2014; Moriguchi et al., 2004; Stehlik et al., 2004). The major BM components in the BBB (i.e., laminin-211, collagen IV) and the prion protein were identified as extracellular ligands for the N-terminal fragment of GPR126. These trigger cAMP signaling, to induce biological effects in Schwann cells through Gαs coupling (Kuffer et al., 2016; Paavola et al., 2014; Petersen et al., 2015). Also, it has been shown that the C-terminal fragment of GPR126 contains a binding site for progesterone and 17-hydroxyprogesterone that triggers its downstream Gαi signaling (An et al., 2022).

Furthermore, GPR126 has been reported to have critical roles in the normal development of diverse organs and tissues, including Schwann cells and peripheral nervous system myelination, bone, inner ear, and placenta (Baxendale et al., 2021). Of note, human GPR126 mutations are linked to several nonneurological diseases, such as adolescent idiopathic scoliosis (Kou et al., 2013), arthrogryposis multiplex congenital (Ravenscroft et al., 2015), and carcinogenesis (An et al., 2022). Although GPR126 has been shown to have a key role in both physiological and pathological angiogenesis through regulation of the expression and activity of VEGFR2, the function of GPR126 at the BBB remains elusive (Cui et al., 2014).

In the present study, we identify Gpr126 as a regulator of BBB establishment and in ischemic stroke, a pathological condition in which the BBB integrity is acutely compromised. We show that Gpr126 is expressed in the CNS endothelium during BBB development, but is reduced in adult quiescent ECs. Endothelial-specific inactivation of Gpr126 in mice (Gpr126*^iECKO^* mice) results in vessel enlargement and vascular abnormalities, which increase neurovascular leakage. RNA sequencing analysis of brain ECs showed that Gpr126 expression is essential to regulate the transcriptional programs for correct BBB development. Indeed, in Gpr126*^iECKO^*mice, TJ organization, BM protein deposition, and pericyte coverage were impaired. We further demonstrate that the loss of BBB characteristics appears to be due to defective angiogenesis. In developing brain ECs and in the retina, Gpr126*^iECKO^* mice showed defects in sprouting, migration, and proliferation during angiogenesis. In unraveling the molecular mechanisms of angiogenesis, we identified Lrp1 as a novel target of Gpr126. Also, Gpr126 binds to collagen IV and increases Lrp1 expression. Gpr126 interacts with Lrp1 and α3β1-integrin, which supports the involvement of the Gpr126-Lrp1 complex in cell migration and in balancing the levels of β1-integrin activation and recycling. We thus identify Gpr126 as a key molecule that positively regulates angiogenesis and BBB development in early postnatal brain. Finally, in adult mice with an established BBB, we demonstrate an unexpected protective effect of Gpr126 deficiency on acute ischemic stroke lesion burden and neurological dysfunction, through dampening of microglia activation.

Collectively, these data provide the first evidence that Gpr126 directly regulates CNS blood vessel morphogenesis and BBB integrity in health. Conversely, an absence Gpr126 is protective during acute breakdown of the mature BBB. Our data suggest that targeting Gpr126 will be a promising therapeutic approach to treat ischemic stroke.

## Results

### Gpr126 is a target of Wnt/β-catenin signaling in the brain microvasculature

Wnt/β-catenin signaling has emerged as a fundamental determinant for acquisition of the specialized phenotype of brain ECs that results in the establishment of the BBB (Daneman et al., 2009; Liebner et al., 2008; Stenman et al., 2008). This suggests that the molecular targets of Wnt/β-catenin signaling have a pivotal role in BBB development. To test this hypothesis, we compared the gene expression profiles of primary murine ECs isolated from the brain microvasculature (cBECs) not treated or treated with the morphogen Wnt3a, a trigger of the canonical Wnt pathway (for 5 days). We identified GPCR 126 (*Gpr126*) as one of the most Wnt3a up-regulated genes (4.8-fold, compared to control cells; p <0.05) among several established Wnt/β-catenin targets, such as axis inhibition protein 2 (*Axin2*), adenomatosis polyposis down-regulated 1 (*Apcdd1*), solute carrier organic anion transporter family member 1C1 (*Slco1c1*), forkhead box F2 (*Foxf2*), forkhead box Q1 (*Foxq1*), zic family member 3 (*Zic3*) and plasmalemma vesicle-associated protein (*Plva*p) (Daneman et al., 2009; Hupe et al., 2017; Liebner et al., 2008; Mazzoni et al., 2017; Sabbagh et al., 2018; Stenman et al., 2008; Zhou et al., 2014) (Figure 1A). No role has been described for *Gpr126* in the development of the BBB to date. Using real-time quantitative polymerase chain reaction (RT-qPCR), we confirmed that *Gpr126* mRNA expression is significantly increased in Wnt3a-stimulated cBECs, with *Axin2* expression as the positive control (Figure 1B). Using not-CNS primary murine ECs isolated from the lung [cLECs, an endothelium that expresses high levels of Gpr126 under untreated conditions (Musa et al., 2019)], we observed that Gpr126 mRNA expression did not change upon Wnt3a stimulation (Figure 1B). Activation of the noncanonical Wnt signaling pathway in cBECs with purified Wnt5a did not increase *Gpr126* mRNA levels, although expression of the selected Wnt5a target gene signal transducer and activator of transcription 2 (*Stat2*) was decreased (Supplementary Figure 1A) (Korn et al., 2014). To the best of our knowledge, these data show for the first time that Gpr126 is a target of canonical Wnt/β-catenin signaling in the mouse brain microvasculature.

**Figure 1.**
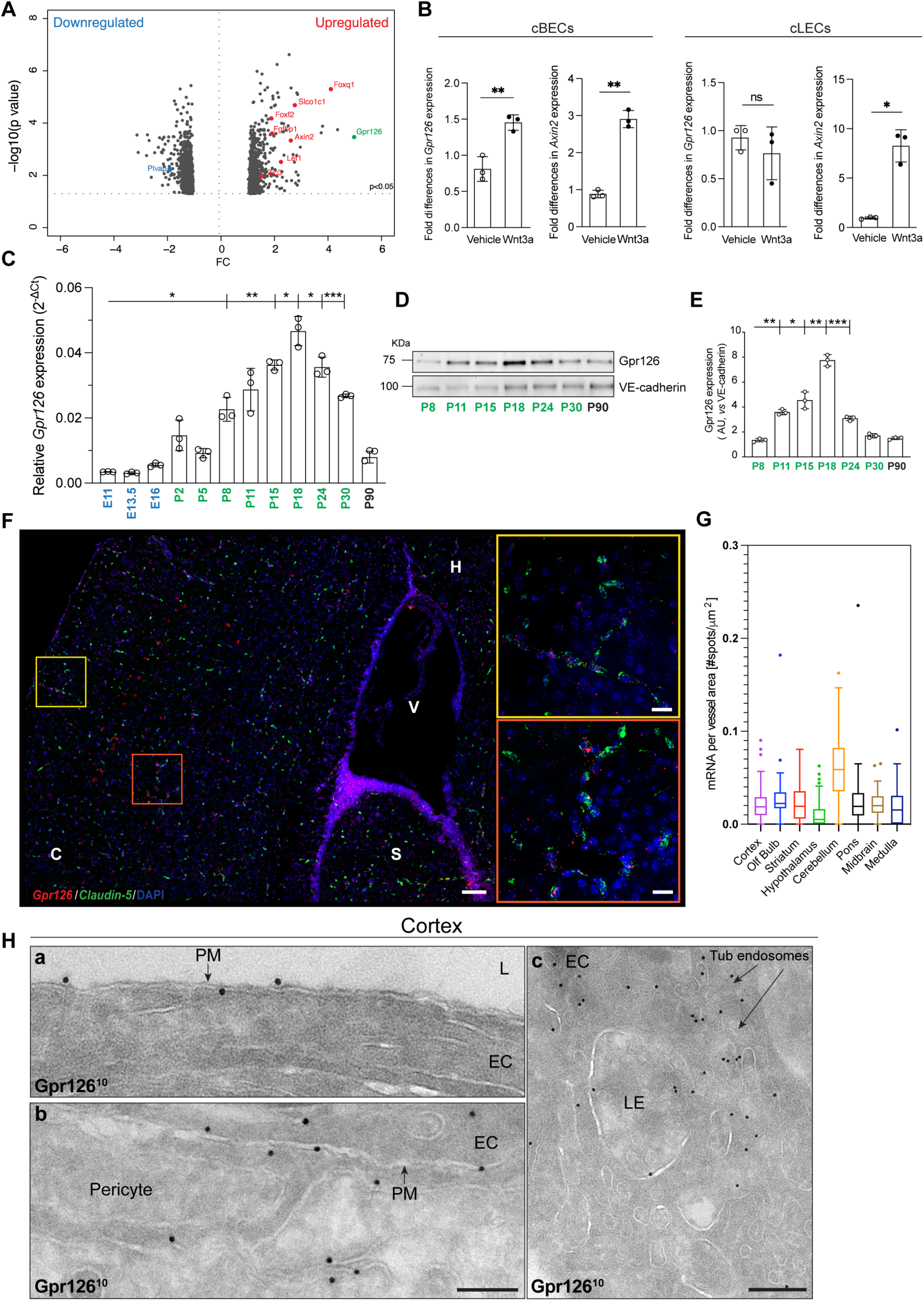
Gpr126 is a novel target of the canonical Wnt/β-catenin signaling pathway and it is expressed in brain vasculature during blood-brain barrier development. **A,** Volcano plot showing transcriptional changes in cultured murine endothelial cells derived from the brain microvasculature (cBECs) in Wnt3a-conditioned medium as treated *versus* control medium. Each dot represents a single gene. Only genes with *P*<0.05 were plotted (Fisher’s least significant difference test). Red dots, up-regulated gene targets of Wnt/β-catenin; blue dots, down-regulated gene targets. *Gpr126* is shown in green. **B,** Quantification of fold differences in *Gpr126* and *Axin2* gene expression in cultured brain-derived (cBECs) and lung-derived (cLECs) endothelial cells from adult WT mice following RT-qPCR analysis. The cells were treated with recombinant Wnt3a or vehicle as control. Each symbol represents a single experiment (n=4 WT mice for each independent experiment, as means ±SD). \**P*<0.05; \*\**P*<0.005; ns=not significant (unpaired t-tests with Welch’s correction). **C,** Quantification of relative *Gpr126* expression in freshly isolated (f)BECs from WT mice at different stages during embryonic (E11-E16) and postnatal (P2-P30) development, and in the adult (P90) following RT-qPCR analysis. Each symbol represents a single experiment (n=8 mice for embryos, n=5 for P2-P5, and n=3 for P8-P90 for each independent experiment, as means ±SD). \**P*<0.05; \*\**P*<0.005; \*\*\**P*<0.0005 (Brown-Forsythe and Welch ANOVA, Dunnett’s T3 multiple comparisons tests). **D,** Representative immunoblotting for Gpr126 in fBECs from WT mice at different postnatal stages (P8-P30) and in the adult (P90). VE-cadherin is shown as loading control. **E,** Gpr126/Ve-cadherin ratios quantified by densitometry scanning and expressed as arbitrary units (AUs), as shown in D. Each symbol represents a single experiment (n=3 mice for each independent experiment, as means ±SD). \**P*<0.05; \*\**P*<0.005; \*\*\**P*<0.0005 (Brown-Forsythe and Welch ANOVA, Dunnett’s T3 multiple comparisons tests). **F,** Representative confocal image of RNA scope *in-situ* hybridization for *Gpr126* (red) and *Claudin-5* (green) mRNA in the cortex of mouse brain at P18. DAPI (4′,6-diamidino-2-phenylindole; blue) stains nuclei. Scale bar: 100 μm. Yellow and orange boxes show magnified cortex regions. Scale bar: 20 μm. C, cortex; S, striatum; V, ventricle; H, hippocampus. **G,** Quantification of *Gpr126* mRNA per vessel area (#spots/μm^2^) in the different brain regions. Each symbol represents a single field in the brain region (3-4 fields/region, n=4 WT mice). **H,** Electron microscopy of Gpr126 immunogold-labeled (10 nm gold, Gpr126^10^) representative cryo-sections of brain capillaries from WT mouse cortex at P18. Black dots, Gpr126 protein in **a**) the luminal side of the plasma membrane of an EC, **b**) the abluminal side of the plasma membrane of an EC and a pericyte, **c**) the tubular endosomes and late endosome in an endothelial cell. EC, endothelial cell, L, lumen, PM, plasma membrane, LE, late endosome, Tub, tubular. Scale bars: 200 nm.

Then, we speculated that Gpr126 has a critical role in the brain endothelium and in BBB specialization, as Wnt signaling is required to establish a functional BBB during embryogenesis and early postnatal life, while in the adult it is reduced to a low level (Liebner et al., 2008). To investigate the involvement of Gpr126 in the physiological establishment of the BBB *in vivo*, we compared the expression of Gpr126 in freshly isolated BECs (fBECs) from mice at different embryonic (E11-16) and postnatal (P2-30) stages and in the adult (P90).

The purities of the fBEC cell preparations were tested for specific markers by RT-qPCR (Supplementary Figure 1B). fBECs showed strong enrichment of the EC-specific marker *Pecam-1*, with low to barely detectable contamination of other cell types of the NVU, such as pericytes (specific marker, *Cspg4*), smooth muscle cells (specific marker, *Acta2*) and astrocytes (specific marker, *Aqp4*) (Supplementary Figure 1B). *Gpr126* transcript levels in the fBECs were low in the embryo (from E11 to E16), and progressively increased in early postnatal life (from P2 to P18), during the formation of the functional BBB. Conversely, in early postnatal life (P24 to P30), *Gpr126* transcript levels decreased, and were highly down-regulated in the adult brain (P90), when the BBB was fully established (Figure 1C). Interestingly, *Gpr126* mRNA levels from E11 to P90 showed significant correlation with the expression of *Axin2* and *β-catenin* (Pearson correlation coefficient, r=0.67, p=0.018, and r=0.69, p=0.012, respectively) which confirmed that *Gpr126* expression was triggered by Wnt/β-catenin signaling in brain ECs (Supplementary Figure 1C, D).

Maturation of the BBB was further confirmed by *Cldn-5* up-regulation (as a specific marker of BBB TJs; (Nitta et al., 2003; Wolburg et al., 2003) and *Plvap* down-regulation (as a component of endothelial fenestrae and caveolae; (Hallmann et al., 1995) (Supplementary Figure 1E, F). Moreover, the Gpr126 protein expression in fBECs from mouse pups at different postnatal stages (from P8 to P30) and adult mice (P90) correlated with *Gpr126* mRNA transcript levels (Figure 1D, E).

Finally, we analyzed Gpr126 receptor localization in the brain microvasculature by RNAscope *in-situ* hybridization, taking advantage of a probe that targets exon 3 of the *Gpr126* gene, which is deleted upon gene inactivation. Thus, we used immortalized brain ECs (iBECs) isolated from this mouse model to validate the probe. We compared *Gpr126* mRNA expression in iBECs without and with a stably expressed murine Gpr126-myc tagged protein (WT, Gpr126-myc) and after *Gpr126* gene inactivation (Gpr126-KO). Confocal images revealed a positive signal in iBECs expressing Gpr126 and claudin-5 (used as an endothelial marker), while no Gpr126 signal was seen for the Gpr126-KO iBECs (Supplementary Figure 1G). Consistent with this, *Gpr126* mRNA levels detected by RT-qPCR showed a similar trend (Supplementary Figure 1H). Hence, we confirmed *Gpr126* expression in blood vessels of the cortex (Figure 1F, G), olfactory bulb, striatum, hypothalamus, cerebellum, pons, midbrain, and medulla (Figure 1G and Supplementary Figure 1I) at P18 (peak of physiological expression of Gpr126, Figure 1C-E).

Electron microscopy analysis of the brain cortex indicated that Gpr126 localized mainly to the plasma membrane of ECs (Figure 1H, panel a) and pericytes (Figure 1H, panel b), which suggested that Gpr126 shows a specific, but non-exclusive, endothelial distribution. In addition, Gpr126 was detected in early endosomes close to the plasma membrane, and in late endosomes and tubular endosomal networks localized within the perinuclear zone of ECs (Figure 1H, panel c).

Thus, we demonstrated that murine Gpr126 expression increases progressively postnatally during the establishment of the BBB and it is regulated by canonical Wnt/β-catenin signaling.

### Gpr126 is required for correct brain vasculature development and BBB function

Given that Gpr126 expression increased postnatally, we investigated the role of Gpr126 in BBB development after birth. To bypass the embryonic lethality of constitutive *Gpr126* deletion due to cardiac defects (Waller-Evans et al., 2010), we generated an inducible endothelial-specific *Gpr126* knockout mouse strain (*Gpr126^iECKO^*) by crossing *Gpr126^flox/flox^*mice with a *Cdh5(PAC)-CreER^T2^* mouse strain (Wang et al., 2010) (Figure 2A). *Gpr126* deletion in mouse ECs was induced by tamoxifen injection, from P1 to P4, and then we analyzed both the recombination efficiency of *Gpr126* and the vascular phenotype at P18 (Figure 2A, B). Gpr126 expression in fBECs from *Gpr126^iECKO^* mice was reduced by 80%, compared to WT by RT-qPCR (Figure 2C). *Gpr126* inactivation led to vasculature alterations in the brain (cortex and striatum) and induced a sparser network of enlarged vessels, vascular abnormalities, and numerous angiogenic sprouts (Figure 2D-F). Consistent with this, in the retina of *Gpr126^iECKO^* mice at P18, arteries and veins appeared more dilated, and contained multiple neovascular tufts, compared to WT (Figure 2G-J). Moreover, Gpr126 reduction decreased the retinal surface vasculature coverage, compared to WT, where the vessel network was fully formed up to the retinal periphery, which suggested that Gpr126 expression is required for appropriate development of the vasculature (Figure 2G). We therefore decided to further explore this process.

**Figure 2.**
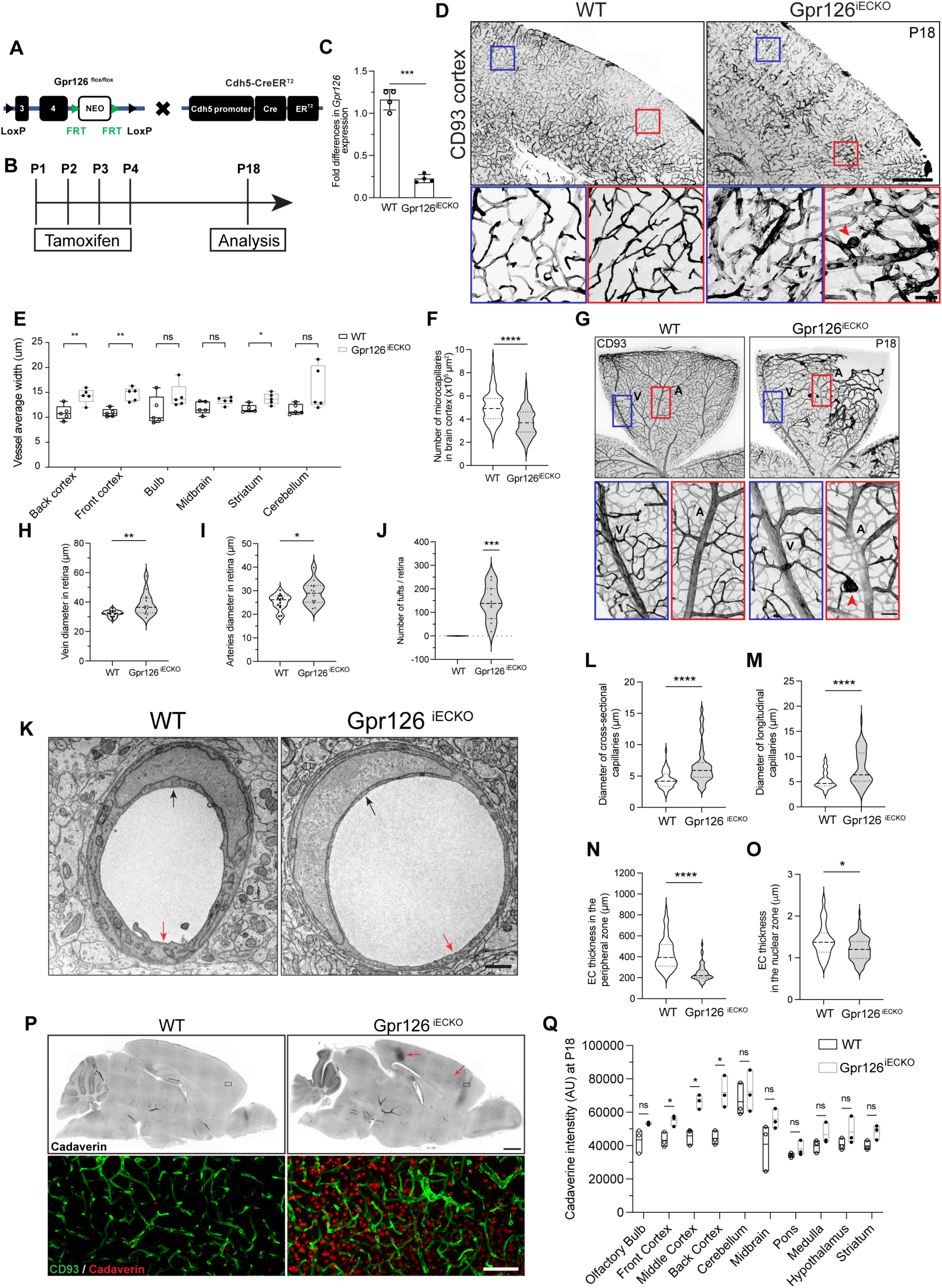
The endothelial Gpr126 is required for formation of the correct brain and retina vasculature. **A,** Generation of *Cdh5-CreER^T2^/Gpr126^flox/flox^* mice. Exons 3 and 4 of the *Gpr126* gene were floxed with LoxP sequences. A neomycin (NEO) cassette flanked by FRT sites was included in the floxed *Gpr126* sequence. *Gpr126*-floxed mice were crossed with mice that expressed Cre recombinase enzyme (Cre) fused with estrogen receptor T2 (ER^T2^) under the control of the *Cdh5* promoter (Cdh5-CreER^T2^). **B,** Experimental protocol for induction of tamoxifen-inducible Cre recombination (P1-P4) and following analysis (P18). **C,** Fold differences in *Gpr126* mRNA in freshly isolated brain endothelial cells from tamoxifen-treated WT and Gpr126^iECKO^ mice at P18. Each symbol represents a mouse (n=4 WT and 4 Gpr126^iECKO^ mice, as means ±SD). ***, *P* ≤0.0005 (t-tests with Welch’s correction). **D,** Representative confocal images for CD93 in vibratome brain sections (100 μm) from WT and Gpr126^iECKO^ mice at P18. Scale bar: 500 μm. Bottom: Red and blue boxes as magnified cortex regions; red arrowhead, a tuft malformation in the vasculature of Gpr126^iECKO^ mouse cortex. Scale bar: 50 μm. **E,** Quantification of mean vessel width in different brain regions of WT and Gpr126^iECKO^ mice at P18. Each symbol represents a mouse (n=5 WT and 5 Gpr126^iECKO^ mice, as means ±SD). *, *P* ≤0.05; **, *P* ≤0.005; ns=not significant (unpaired t-tests with Welch’s correction for all datasets except from bulb data that were analyzed by Mann Whitney tests). **F,** Quantification of the number of microcapillaries in the cortex of WT and Gpr126^iECKO^ mice at P18 per area (n=5 WT and 5 Gpr126^iECKO^ mice, as means ±SD). ****, *P* ≤0.00005 (unpaired t-tests with Welch’s correction). **G,** Representative confocal images for CD93 in the retina of WT and Gpr126^iECKO^ mice at P18. Scale bar: 500 μm. Bottom: Blue and red rectangles, magnified retina regions; red arrowhead, a tuft malformation in the retinal vasculature of Gpr126^iECKO^ mouse. V, vein, A, artery. Scale bar: 200 μm. **H, I,** Quantification of vein **(H)** and artery **(I)** diameters in retina of WT and Gpr126^iECKO^ mice at P18. Each symbol represents a single retina (n=7 WT and 7 Gpr126^iECKO^ mice, as means ±SD). *, *P* ≤0.05; **, *P* ≤0.005 (unpaired t-tests with Welch’s correction). **J**, Quantification of the number of tuft malformations in retina of WT and Gpr126^iECKO^ mice at P18. Each symbol represents a single retina. (n=6 WT and 6 Gpr126^iECKO^ mice, as means ±SD). ***, *P* ≤0.0005 (Wicoxon signed rank tests). **K,** Representative electron microscopy of cross sections of blood capillaries in WT and Gpr126^iECKO^ mice. Images were acquired at the same magnification. Black arrows, luminal perinuclear zone; red arrows, peripheral zone of the EC. Scale bar: 1 μm. **L-O,** Quantification of cross-sectional capillary **(L)** and longitudinal capillary **(M)** diameters, and EC thickness in the peripheral zone **(N)** and in the nuclear zone **(O)** (n=3 WT and 3 Gpr126^iECKO^ mice, as means ±SD). *, *P* ≤0.05; ****, *P* ≤0.00005 (unpaired t-tests with Welch’s correction). **P,** Representative confocal images for cadaverin leakage in vibratome brain sections (100 μm) from WT and Gpr126^iECKO^ mice at P18. Red arrows, area of leakage in the cortex. Scale bar: 1 mm. Bottom: Magnified cortex area of the sections stained for CD93 (green) and cadaverin (red). Scale bar: 100 μm. **Q,** Quantification of cadaverine leakage in different brain regions of WT and Gpr126^iECKO^ mice at P18, as shown in **P**. Each symbol represents a mouse (n=3 WT and 3 Gpr126^iECKO^ mice, as means ±SD). *, *P* ≤0.05; unpaired t-tests with Welch’s correction).

Here, we used electron microscopy to measure the vasculature in the brain cortex of WT and *Gpr126^iECKO^* mice. The capillary diameters significantly increased (Figure 2K-M), while the thickness of the ECs decreased in both peripheral (Figure 2N) and nuclear (Figure 2O) zones of *Gpr126^iECKO^*mice, compared to WT.

To investigate whether the absence of Gpr126 affects BBB properties by aberrant vascular morphogenesis, we analyzed the expression of Plvap, a marker for immature and not completely developed vasculature of the BBB (Wisniewska-Kruk et al., 2016). The Plvap protein and mRNA expression levels were significantly increased in the cortex vasculature of *Gpr126^iECKO^* mice, compared to WT littermates, which suggested that Gpr126 is involved in regulation of BBB maturation and development (Supplementary Figure 2A, B). Consistent with this, the BBB permeability was significantly increased. As shown in Figure 2P, leakage of the low molecular weight tracer cadaverine was increased in the cortex area, where the vessels were most enlarged, which suggested that the BBB function was compromised in *Gpr126^iECKO^* mice (Figure 2P, Q).

### Gpr126 stabilizes junctions, and promotes BM protein deposition, and endothelial cell– pericyte interactions

The BBB paracellular permeability is determined by the junctional complexes that tightly closing the brain ECs, as TJs, adherens junctions, and gap junctions (Stamatovic et al., 2016). To study whether changes in Gpr126 expression altered the junction assembly and function, and thus affected the paracellular permeability, we analyzed the gene expression levels of the components of TJs (*F11 receptor* [*F11r*], *Cldn5, Tight junction protein 1* [*Tjp1*], *Occludin* [*Ocln*]) and adherens junctions (*Junction plakoglobin* [*Jup*], *Cadherin 5* [*Cdh5*], *Pecam-1*) in fBECs. The mRNA levels of *Cldn5*, *F11r,* and *Cdh5* were significantly up-regulated in fBECs from *Gpr126^iECKO^* mice, compared to WT, whereas *Tjp1*, *Ocln,* and *Jup* mRNA levels were not affected (Supplementary Figure 2C). The increase in claudin-5 and junctional adhesion molecule-A expression was also confirmed at the protein level in the vasculature of the cortex of *Gpr126^iECKO^* mice, compared to WT, by immunofluorescence (Supplementary Figure 2D-F).

To quantify differences in junction shapes between WT and *Gpr126^iECKO^* brain vessels, the open-source “Patching” image analysis software was used (Bentley et al., 2014). This is a tool that was previously validated to ensure unbiased hand classification of differential remodeling/stable endothelial junction shapes. Here, there were significantly more “strongly remodeling” junctions (with a serrated shape) and significantly fewer “strongly stable” junctions (with a straight shape) in *Gpr126^iECKO^* brain ECs, compared to WT, which suggested a positive role of Gpr126 in junction stabilization (Supplementary Figure 2G). The automatic feature detection quantification of this software demonstrated that hand classification was consistent, as patches that were classed as strongly remodeling contained more strongly serrated large (area, >50 pixels) objects than strongly stable classes, where objects scored higher on the automated straightness scores (Supplementary Figure 2H, I) independent of whether the data was from WT or *Gpr126^iECKO^* brains.

The BBB extracellular matrix (ECM) components are also crucial for BBB establishment, integrity and function. Brain ECs are embedded in the BM with pericytes, and both cell types contribute to the secretion of components like collagen IV and fibronectin, as well as laminin-211 (α2β1γ1), laminin-511 (α5β1γ1), and laminin-411 (α4β1γ1) (Nirwane and Yao, 2018). As an adhesion GPCR, Gpr126 has an important role during development via many processes, although primarily through cell–cell and cell–ECM interactions, by binding to collagen IV or laminin-211. Thus, we analyzed the structure and composition of the vascular BM coupled to pericytes, using coverage quantification by imaging. At the ultrastructural level the brain cortex blood capillaries and the retina BM of *Gpr126^iECKO^*mice showed irregular thickness, which was characterized by zones without protein matrix, compared to WT mice, where the BM appeared as a continuous layer (Fig. 3A, Supplementary Figure 3A).

**Figure 3.**
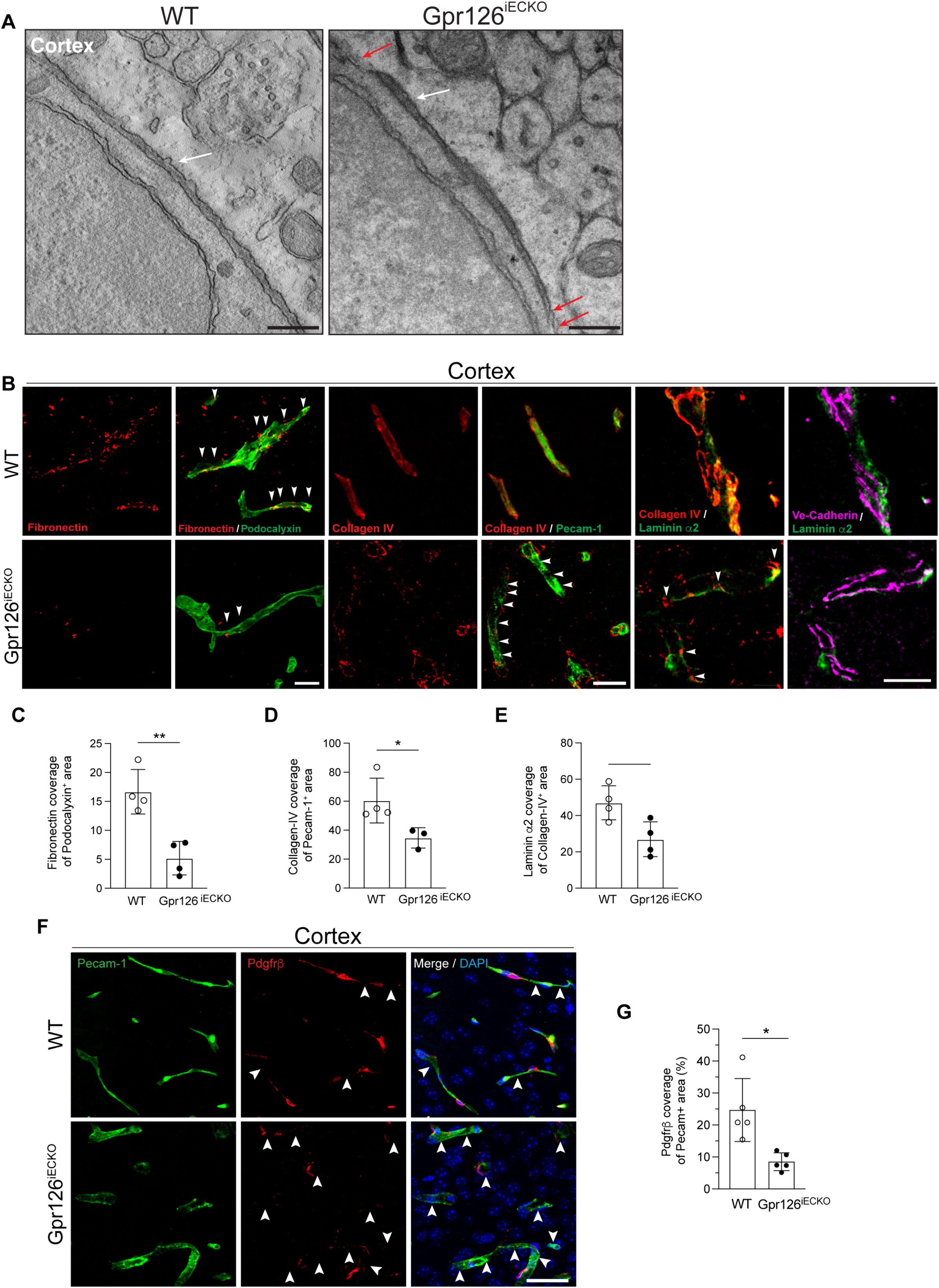
Endothelial Gpr126 is required for correct basement membrane (BM) formation and pericyte vessel coverage. **A,** Representative tomography virtual slices of brain blood capillary longitudinal sections from WT and Gpr126^iECKO^ mice at P18. White arrows, regular thickness of the BM; red arrows, areas of interrupted or thin BM in the ECs. See Supplementary Figure 3A for full image field. Scale bar: 350 nm. **B,** Representative confocal images of brain cortex cryosections (4 μm) from WT and Gpr126^iECKO^ mice at P18. For the vasculature, podocalyxin (green), or Pecam-1 (green), or Ve-cadherin (magenta) are shown. The BM was stained for fibronectin (red), collagen IV (red), or laminin α2 (green). Fibronectin/Podocalyxin: arrowheads show colocalization of the two proteins. Collagen IV/Pecam-1: arrowheads show discontinuous collagen IV staining. Collagen IV/Laminin α2: arrowheads show absence of colocalization of the two proteins. See Supplementary Figure 3B for full image field. Scale bar: 100 μm. **C-E**, Quantification of fibronectin coverage of podocalyxin-positive areas (**C**), collagen IV coverage of Pecam-1-positive areas (**D**), and collagen IV coverage of laminin α2-positive areas (**E**), as shown in **B**. Each symbol represents a mouse (n=4 WT and 3 or 4 Gpr126^iECKO^ mice, as means ±SD). *, *P* ≤0.05; **, *P* ≤0.005 (unpaired t-tests with Welch’s correction). **F,** Representative confocal images for Pecam-1 (ECs, green) and Pdgfrβ (pericytes, red) in brain cortex cryosections (4 μm) of WT and Gpr126^iECKO^ mice at P18. DAPI stains nuclei. Arrowheads indicate discontinuous or absent Pdgfrβ staining. Scale bar: 200 μm. **G,** Quantification of Pdgfrβ coverage of Pecam-1-positive areas, as shown in **F**. (n=5 WT and 5 Gpr126^iECKO^ mice, as means ±SD). *, *P* ≤0.05 (unpaired t-tests with Welch’s correction).

We then investigated the BM components, fibronectin, collagen IV and laminin α2, expression in the brain cortex blood capillaries by immunofluorescence (Figure 3B). Deposition of fibronectin around the podocalyxin-positive blood vessels of *Gpr126^iECKO^* mice was significantly reduced throughout the cortex (Figure 3B, C and Supplementary Figure 3B). Collagen IV showed integral and continuous structures around the Pecam-1-positive microvessels in the WT cortex that were lost in *Gpr126^iECKO^*mice, where collagen IV deposition was discontinuous, irregular, and markedly different from WT (Figure 3B, D). Immunoreactivity quantification revealed that in the absence of Gpr126, the collagen IV endothelial surface coverage decreased (66%), compared to WT (Figure 3D). Moreover, collagen IV and laminin α2 co-distribution in the BM of vascular structures was significantly reduced by 45% in the microvasculature of *Gpr126^iECKO^*mice, compared to WT (Figure 3B, E). The mislocalization of collagen IV and laminin α2 was confirmed in cBECs, which suggested that secretion and aberrant deposition of laminin α2 is specifically mediated by the inactivation of Gpr126 in ECs (Supplementary Figure 3C). To address whether altered fibronectin and collagen IV deposition in *Gpr126^iECKO^* mice was due to reduced expression of *Fn1*, *Col4a1*, and *Col4a2*, we performed RT-qPCR in fBECs of WT and *Gpr126^iECKO^*pups at P18. *Fn1* mRNA levels were comparable between WT and these mutant mice, while *Col4a1* and *Col4a2* were slightly increased (Supplementary Figure 3D-F), which suggested that collagen IV reduction around the vasculature was not caused by decreased mRNA expression in ECs, but more likely by impaired deposition of the mature protein into the vascular BM. Consistent with this, there were increased transcript levels of the metalloproteases *Mmp3*, *Mmp9*, and *Mmp14* in ECs of *Gpr126^iECKO^* mice, compared to WT (Supplementary Figure 3G-I, which might cause BBB disruption through ECM degradation and subsequent pericyte loss around the microvasculature. Indeed, brain capillary dual immunostaining for Pdgfr-β (a pericyte marker) and Pecam-1 (an EC marker) showed that pericyte coverage was significantly decreased by 60% in *Gpr126^iECKO^*, compared to WT (Figure 3F, G).

Similar results were obtained for the retina vasculature. Control mice blood vessels were positive for CD13, a surface marker for pericytes, while *Gpr126^iECKO^* mice had a significantly reduced number of pericytes, with abnormal blood vessels with an irregular and disorganized distribution of pericytes, compared to WT mice (Figure 4A, B). In line with these findings, α-smooth muscle actin (αSMA) immunoreactivity was detected only in arteries, and it was significantly reduced in *Gpr126^iECKO^* mice (Supplementary Figure 4 A, B).

**Figure 4.**
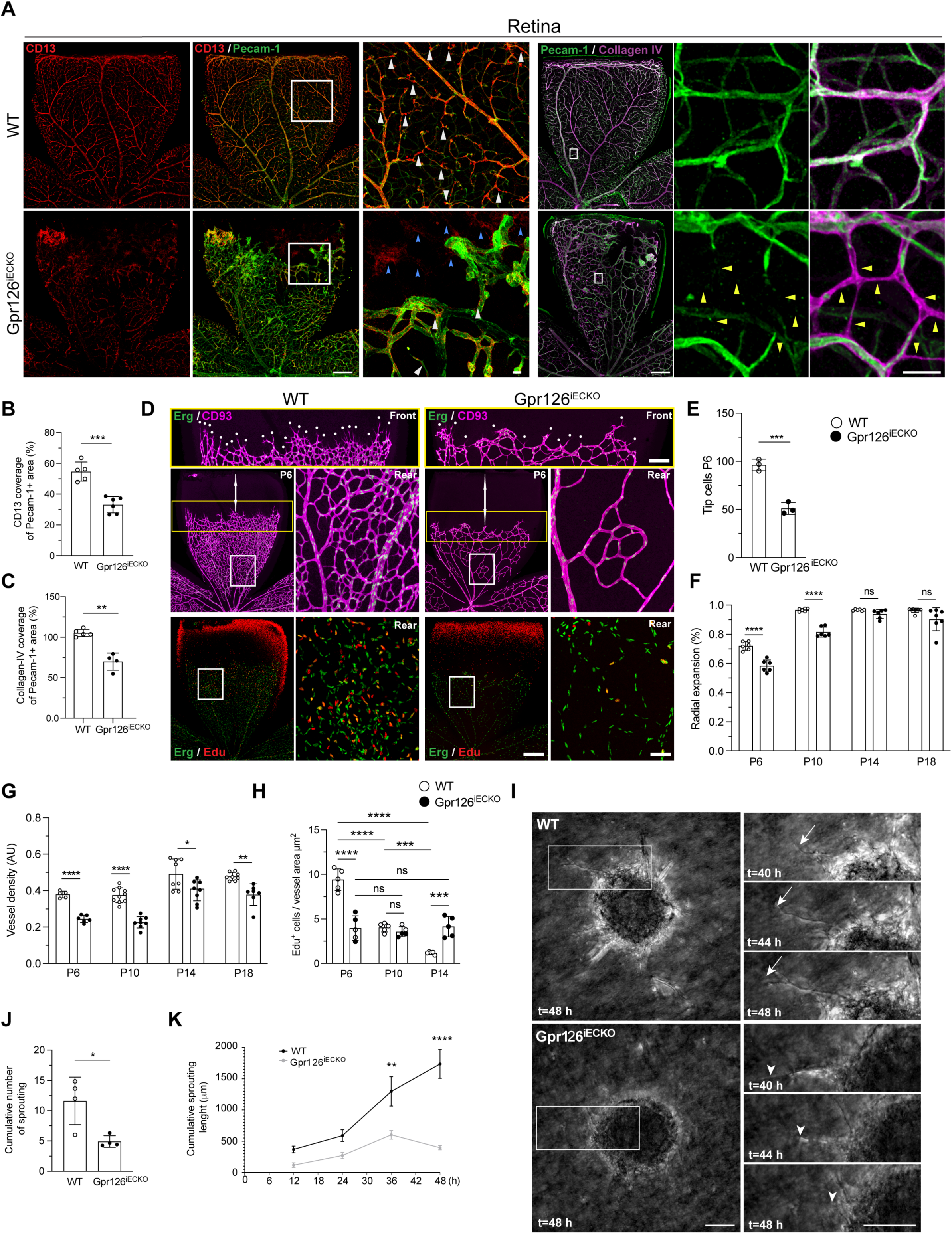
Endothelial Gpr126 is required for angiogenesis in the retina and in the brain. **A,** Representative confocal images for Pecam-1 (ECs, green) CD13 (pericytes, red) and collagen IV (BM, magenta) in WT and Gpr126^iECKO^ mouse retinas at P18. Scale bars: 500 μm. White squares and rectangles are shown magnified on the right for each image. Scale bars: 50 μm. White arrowheads in CD13/Pecam-1 staining show most of the pericyte bodies; blue arrowheads show CD13 staining without Pecam-1 staining (retraction); white arrowheads in Pecam-1/collagen IV staining show collagen IV vessel-like structures without Pecam-1 staining (retraction) **B,** Quantification of CD13 coverage of Pecam-1-positive areas, as shown in **A**. Each symbol represents a retina for each mouse (n=5 WT and 6 Gpr126^iECKO^ mice, as means ±SD). ***, *P*≤0.0005 (unpaired t-tests with Welch’s correction). **C,** Quantification of collagen IV coverage of Pecam-1-positive areas, as shown in **A**. Each symbol represents a retina for each mouse (n=5 WT and 4 Gpr126^iECKO^ mice, as means ±SD). **, *P*≤0.005 (unpaired t-tests with Welch’s correction). **D,** Representative confocal images for CD93 (pan-EC membrane and filopodia at P6, magenta), Erg (EC nuclei, green) and Edu (proliferating cell nuclei, red) in WT and Gpr126^iECKO^ mouse retinas at P6. Retina vessels are shown for the full petal, migrating front (Front, yellow rectangles, magnification) and proximal to the optic nerve (Rear, white rectangle, magnification). The bidirectional arrow indicates the distance between the migrating vasculature front and the end of the petal. White dots (in magnifications) indicate tip cells (filopodia-rich cells); Petal, scale bar: 500 μm; rear, scale bar: 100 μm; front, scale bar: 250 μm. **E,** Quantification of the tip cells shown in the yellow rectangle in **D**. Each symbol represents a retina for each mouse. (n=3 WT and 3 Gpr126^iECKO^ mice, as means ±SD). ***, *P*≤0.0005 (unpaired t-tests with Welch’s correction). **F,** Quantification of radial expansion of the retinal vasculature post-natally (P6, P10, P14, P18) in WT and Gpr126^iECKO^ mice. Each symbol represents a retina for each mouse (n=6-8 WT and 6-7 Gpr126^iECKO^ mice, as means ±SD, WT *vs* Gpr126^iECKO^) ****, *P*≤0.00005, ns=not significant (unpaired t-tests with Welch’s correction). **G,** Quantification of vessel density in the retinal vasculature post-natally (P6, P10, P14, P18) for WT and Gpr126iECKO mice. Each symbol represents a retina for each mouse. (n=5-10 WT and 6-9 Gpr126^iECKO^ mice, as means ±SD, WT *vs* Gpr126^iECKO^). *, *P*≤0.05; **, *P*≤0.005; ****, *P*≤0.00005 (unpaired t-tests with Welch’s correction). **H,** Quantification of Edu-positive ECs per μm^2^ of vessel area post-natally (P6, P10, P14, P14) in WT and Gpr126^iECKO^ mouse retinas. Each symbol represents a retina for each mouse. (n=5 WT and 5 Gpr126^iECKO^ mice, as means ±SD). ***, *P*≤0.0005, ****, *P*≤0.00005, ns=not significant (2-way ANOVA). **I,** Representative phase-contrast images of sprouting spheroids from fBECs from WT and Gpr126^iECKO^ mice at P18, after 48 h stimulation with VEGF (80 ng/mL) and Fgfb (50 ng/mL). Right: Rectangular boxes show magnified representative phase-contrast images of the sprouting spheroids at different times (t=40 h, 44 h and 48 h) (See Supplementary Movie 1 for time-lapse images from the assay). Arrows show a sprouting EC over time. Arrowheads show a retracting EC over time. Scale bar: 40 μm. **J,** Quantification of the cumulative numbers of sprouts per spheroid of WT (n = 16 spheroids) and Gpr126^iECKO^ (n = 16 spheroids) fBECs. Each symbol represents a single experiment (n=4 WT and 4 Gpr126^iECKO^ mice for each experiment, as means ±SD). *, *P*≤0.05 (t-tests with Welch’s correction). **K,** Quantification of cumulative sprouting lengths of WT (n = 12 spheroids) and Gpr126^iECKO^ (n = 12 spheroids) fBECs, after 12, 24, 36 and 48 h (n=6 WT and 6 Gpr126^iECKO^ mice for each experiment, as means ±SD). **, *P*≤0.005; ****, *P*≤0.00005 (2-way ANOVA).

Surprisingly, the presence of CD13-positive and Pecam-1–negative structures in the front of the retina of *Gpr126^iECKO^* mice at P18 implied that the vessels underwent regression (Figure 4A, B). This process was also confirmed by the appearance of numerous collagen IV-positive and Pecam-1–negative stained structures that represented empty matrix sleeves in the *Gpr126^iECKO^*vasculature and decorated blind-ended vessels (Figure 4A, C).

Collectively, these findings indicate that Gpr126 is essential for correct BM deposition, which, in turn, is crucial for guiding brain vessels into the spatially distributed and functional network that is optimally suited to maintain BBB homeostasis (Egginton and Gaffney, 2010).

### Gpr126 induces angiogenesis in the CNS

Brain angiogenesis is very tightly coupled to the formation of the BBB. Thus, we investigated the role of Gpr126 in developmental retinal angiogenesis after birth. At P6, the growing superficial vessel plexus had not yet reached the periphery of the retina, and numerous sprouts were visible at the angiogenic growth front. Tamoxifen-inducible Gpr126 depletion in ECs at P1 led to significantly decreased numbers of tip cells, radial expansion, and vascular density, which suggested that Gpr126 expression is important for CNS and retina angiogenesis (Figure 4D-G).

At P14, the extension of the superficial vascular network was complete in both control (WT) and *Gpr126^iECKO^*mice (Figure 4F and Supplementary Figure 4C). However, the formation of the deep plexus network was still significantly delayed in the *Gpr126^iECKO^* retina (Supplementary Figure 4C), which suggested that the deletion of endothelial Gpr126 impaired the migratory performance of ECs. Moreover, at P14, in the absence of Gpr126, the vessel density remained significantly reduced, compared to WT, and the veins became enlarged (Figure 4G and Supplementary Figure 4C-E).

As an indicator of endothelial proliferation, substantial ethynyl deoxyuridine incorporation (Edu) was seen for WT Erg-positive EC nuclei at P6, which significantly decreased over time to P14, where Edu-labeled ECs were almost exclusively in the most distal, peripheral sections of veins (Figure 4D, H and Supplementary Figure 4F). Conversely, Edu and Erg double-positive cells in the *Gpr126^iECKO^* retina were significantly reduced at P6, compared to WT, and showed similar numbers to P14 (Figure 4D, H and Supplementary Figure 4F). It is of note that at P14, Erg-positive EC nuclei were significantly increased in the *Gpr126^iECKO^* retina, compared to WT, which suggested that the absence of Gpr126 delayed vasculature development (Figure 4H and Supplementary Figure 4F).

To further investigate the involvement of Gpr126 in brain angiogenesis, we performed *ex-vivo* experiments using cBECs of WT and *Gpr126^iECKO^*mice at P18. Cell proliferation and migration are both tightly connected with angiogenesis; therefore, we investigated Gpr126 involvement in cell proliferation and migration. Proliferation was measured using bromodeoxyuridine incorporation and Ki67 staining. In ECs from *Gpr126^iECKO^* mice, proliferation was reduced under standard culture conditions, compared to WT (Supplementary Figure 4G-I). Also, ECs from *Gpr126^iECKO^* mice showed slower cell migration, compared to WT (as closing of the wound in a scratch wound healing assay) (Supplementary Figure 4J, K).

Finally, to determine how Gpr126 regulates EC movement during angiogenesis, multicellular spheroids composed of fBECs from WT and *Gpr126^iECKO^*mice were placed in a three-dimensional collagen matrix to monitor VEGF-Fgfb–induced angiogenesis. Depletion of Gpr126 strongly decreased sprout formation and elongation from spheroids (Figure 4I-M), a further confirmation of the role of Gpr126 in angiogenesis, with a reduction in new blood vessels upon loss of Gpr126 in ECs. Moreover, live-cell time-lapse showed that the sprouting length increased over time in WT ECs (Figure 4I-K, Supplementary Movie 1), while in the absence of Gpr126, the sprout length decreased, with regression events after 2 days of growth (Figure 4I, K). Also, in the retina at P18, the absence of Gpr126 significantly decreased the vascular density, and there was a tendency to a reduction in the radial expansion, which suggested that the formed vessels were not stable and were prone to regression (Figure 4F, G).

Collectively, these data demonstrate that impairment of proliferation and migration in Gpr126-deficient ECs contributes to the defects in angiogenesis in the CNS of *Gpr126^iECKO^* mice.

### Lrp1 is a novel target of Gpr126

To define the Gpr126 mechanism of action during BBB formation, we profiled the brain endothelial transcriptional changes upon Gpr126 depletion at P18. Differential expression analyses showed that *Gpr126^iECKO^* led to a marked rewiring of transcription, compared to WT (Supplementary Figure 5A, B).

According to gene set enrichment analysis, loss of Gpr126 resulted in global down-regulation of a variety of biological processes. Thus, we focused on a subset of gene ontology (GO) terms related to specific BBB properties and functions (Figure 5A), with a negative enrichment score for the majority of these genes (Supplementary Figure 5C). Moreover, the leading-edge genes (i.e., the subset of genes that contributes the most to the signal) of these altered GO terms showed concordant down-regulation among the samples (Figure 5B). Finally, we selected for validation those genes that were present in at least 75% of all of the selected GO terms (Figure 5C and Supplementary Table I). Among these genes, we focused on *lrp1*, as over recent years an increased number of reports have demonstrated that Lrp1 is involved in EC functions, such as BBB transcytosis, permeability, and angiogenesis (Mao et al., 2017; Nakajima et al., 2014; Storck et al., 2016; Zhao et al., 2016). In addition, Lrp1 regulates several TJ proteins and mediates the clearance of major ECM-degrading proteinases (Zhao et al., 2016).

**Figure 5.**
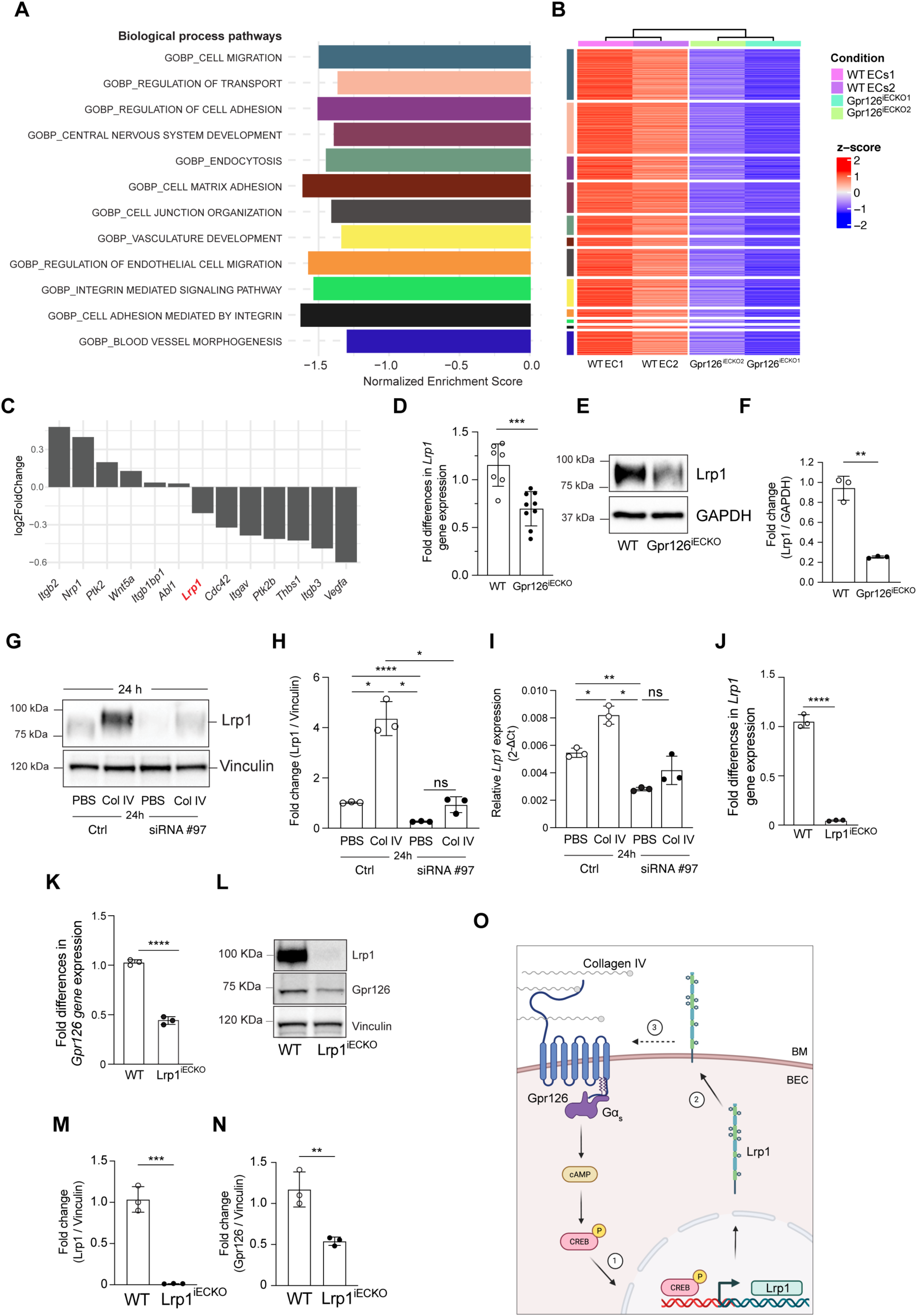
Lrp1 is a novel target of the Gpr126 signaling pathway. **A-C,** fBECs were isolated from WT and Gpr126^iECKO^ mice at P18 and the mRNA was processed for bulk RNA sequencing. **A**, Normalized Enrichment Scores (NES) of selected enriched pathways obtained from gene set enrichment analysis (GSEA) as differentially expressed genes under Gpr126^iECKO^ *vs* WT conditions (pval ≤0.01, padj ≤0.05, at least 30 altered genes). Moderated t-statistics were used to rank the genes. GSEA was performed on the GO Biological process gene sets from the GSEA molecular signatures database. **B,** Heatmap showing the Z-scores of the leading-edge genes, the core genes that account for the enrichment signal obtained from GSEA analysis for selected GO Biological Process gene sets in each of the two biological replicates under each condition (n=4 WT and 4 Gpr126^iECKO^ mice for each replicate). The Z-scores were calculated for each gene starting from the DeSeq2 variance stabilized expression value corrected for batch effects (using the Bioconductor limma package Remove BatchEffect function) and then normalized with the Bioconductor pheatmap package scale_rows function. **C,** Log2 of fold-change values of selected genes differentially expressed in Gpr126^iECKO^ *vs* WT conditions. Differential expression analysis was performed using DeSeq2. *Lrp1* (red) was one of the most common genes which is present in at least 75% of all of the selected GO terms. **D,** Quantification of the fold differences in *Lrp1* gene expression in fBECs from WT and Gpr126^iECKO^ pups at P18. Each symbol represents a mouse (n=7 WT and 9 Gpr126^iECKO^ mice, as means ±SD). ***, *P*<0.0005 (unpaired t-tests with Welch’s correction). **E,** Representative immunoblotting for Lrp1 in fBECs from WT and Gpr126^iECKO^ mice at P18. GAPDH is shown as the loading control. **F**, Lrp1/GAPDH ratio quantified by densitometry scanning and expressed as fold change, as show in **E**. Each symbol represents a single experiment (n=3 WT and 3 Gpr126^iECKO^ mice for each independent experiment, means ±SD). \*\**P*<0.005 (unpaired t-tests with Welch’s correction). **G,** Representative immunoblotting for Lrp1 in cBECs isolated from P18 pups transfected with nontargeting small-interfering RNA (siRNA; Ctrl [control]) or with siRNAs against Gpr126 (#97) and treated with collagen IV (or PBS, as vehicle) for 24 h. Vinculin is shown as a loading control. **H,** Lrp1/Vinculin ratio quantified by densitometry scanning and expressed as fold changes, as shown in **G**. Each symbol represents a single experiment (n=3 WT and 3 Gpr126^iECKO^ mice for each independent experiment, as means ±SD). *, *P*<0.05; ***, *P*<0.0005; ****, *P*<0.00005; ns=not significant (Brown-Forsythe and Welch ANOVA, Dunnett’s T3 multiple comparison tests). **I,** Quantification of relative gene expression of *Lrp1* in cBECs isolated from pups at P18, as shown in **G**. Each symbol represents a single experiment (n=3 WT and 3 Gpr126^iECKO^ mice for each independent experiment, as means ±SD). *, *P*<0.05; **, *P*<0.005; ns=not significant (Brown-Forsythe and Welch ANOVA, Dunnett’s T3 multiple comparison tests). **J,** Quantification of fold differences in gene expression of *Lrp1* in cBECs from adult WT and Lrp1^iECKO^ mice. Each symbol represents a single experiment (n=3 WT and 3 Gpr126^iECKO^ mice for each independent experiment, as means ±SD). ****, *P*<0.00005 (unpaired t-tests with Welch correction). **K,** Quantification of fold differences in gene expression of *Gpr126* in cBECs from adult WT and Lrp1^iECKO^ mice. Each symbol represents a single experiment (n=3 WT and 3 Gpr126^iECKO^ mice for each independent experiment, as means ±SD). ****, *P*<0.00005 (unpaired t-tests with Welch correction). **L,** Representative immunoblot for Lrp1 and Gpr126 in cBECs from adult WT and Lrp1^iECKO^ mice. Vinculin is shown as a loading control. **M,** Lrp1/Vinculin ratio quantified by densitometry scanning and expressed as fold change, as shown in **L**. Each symbol represents a single experiment (n=3 WT and 3 Gpr126^iECKO^ mice for each independent experiment, as means ±SD). ***, *P*<0.0005; unpaired t-tests with Welch’s correction). **N,** Gpr126/Vinculin ratio quantified by densitometry scanning and expressed as fold changes, as shown in **L**. Each symbol represents a single experiment (n=3 WT and 3 Gpr126^iECKO^ mice for each independent experiment, means ±SD). **, *P*<0.005 (unpaired t-tests with Welch’s correction). **O**, Schematic model for Gpr126 regulation of Lrp1 expression and *vice versa*. **1.** Upon collagen IV binding to Gpr126 receptor, the increase of cAMP production phosphorylates Creb, which in turn translocates to the nucleus and increases the transcription of Lrp1; **2.** Thus, Lrp1 localizes at the plasma membrane; and **3.** supports Gpr126 expression and signaling. Dashed arrow indicates that the molecular mechanisms through which Lrp1 regulates Gpr126 expression remain to be defined. BEC, brain endothelial cell; BM, basement membrane.

Using RT-qPCR, we confirmed that both the Lrp1 protein and *lrp1* mRNA are expressed in WT ECs, and they are significantly reduced in the absence of Gpr126 in fBECs from mice at P18 (Figure 5D-F). We then further studied Lrp1 using several approaches to modulate Gpr126 expression and its downstream signaling activity. Acute Gpr126 down-regulation by small-interfering RNAs in WT cBECs (Supplementary Figure 6A-C) led to significant decreases in Lrp1 expression at both the protein and mRNA levels, compared with control cells (Figure 5G-I). Next, we investigated whether Lrp1 expression is triggered by the activation of Gpr126 receptor binding to its known ligand collagen-IV, which produces cyclic adenosine monophosphate (cAMP) and induces phosphorylation of Creb (Paavola et al., 2014). Treatment of WT cBECs with soluble collagen-IV increased Creb phosphorylation at S133, while in the absence of Gpr126 the levels of p-Creb^S133^ were low, and were comparable to vehicle only treated cells (Supplementary Figure 6D, E). Moreover, collagen-IV treatment of WT ECs cells for 24 h stimulated Gpr126 signaling by increasing the Lrp1 protein and mRNA levels, which was not seen upon Gpr126 depletion (Figure 5G-I).

Finally, to better describe the connections between Lrp1 and Gpr126 receptors, we investigated whether the absence of Lrp1 affected the expression of Gpr126, taking advantage of WT and Lrp1^iECKO^ brain ECs isolated from recombined adult Slco1c1-Cre-ER^T2^/Lrp1^flox/flox^ mice (Storck et al., 2016). In the Lrp1^iECKO^ brain ECs, both Lrp1 and Gpr126 mRNA and protein levels were down-regulated, compared to WT (Figure 5J-N).

These data (as summarized in Figure 5O) demonstrate that Lrp1 is a specific target of Gpr126 upon collagen-IV binding in brain ECs, with transcriptional regulation between Lrp1 and Gpr126.

### Gpr126 interacts with Lrp1 and α3β1-integrin to regulate angiogenesis

A role for LRP1 in angiogenesis has emerged in recent years (Mao et al., 2017). It has been shown that Lrp1 mediates β1-integrin interactions and acts as a potential regulator of cell migration and adhesion (Rabiej et al., 2016), and that β1-integrin has an important role in angiogenesis through VEGF signaling (Avraamides et al., 2008).

In line with these data, we observed that some of the down-regulated genes in at least 75% of all of the selected GO terms (Figure 5C) have functions in angiogenesis, like cell division control protein 42 homolog (*Cdc42*), integrin subunit alpha V (*Itgav*), protein tyrosine kinase 2 beta (*Ptk2b*), thrombospondin 1 (*Thbs1*), integrin subunit beta *3* (*Itgb3*), and vascular endothelial growth factor A (*Vegfa*) (Figure 5D and Table 1). Hence, we postulated that Gpr126 positively regulates angiogenesis through the interaction with Lrp1–β1-integrin.

**Table 1.**
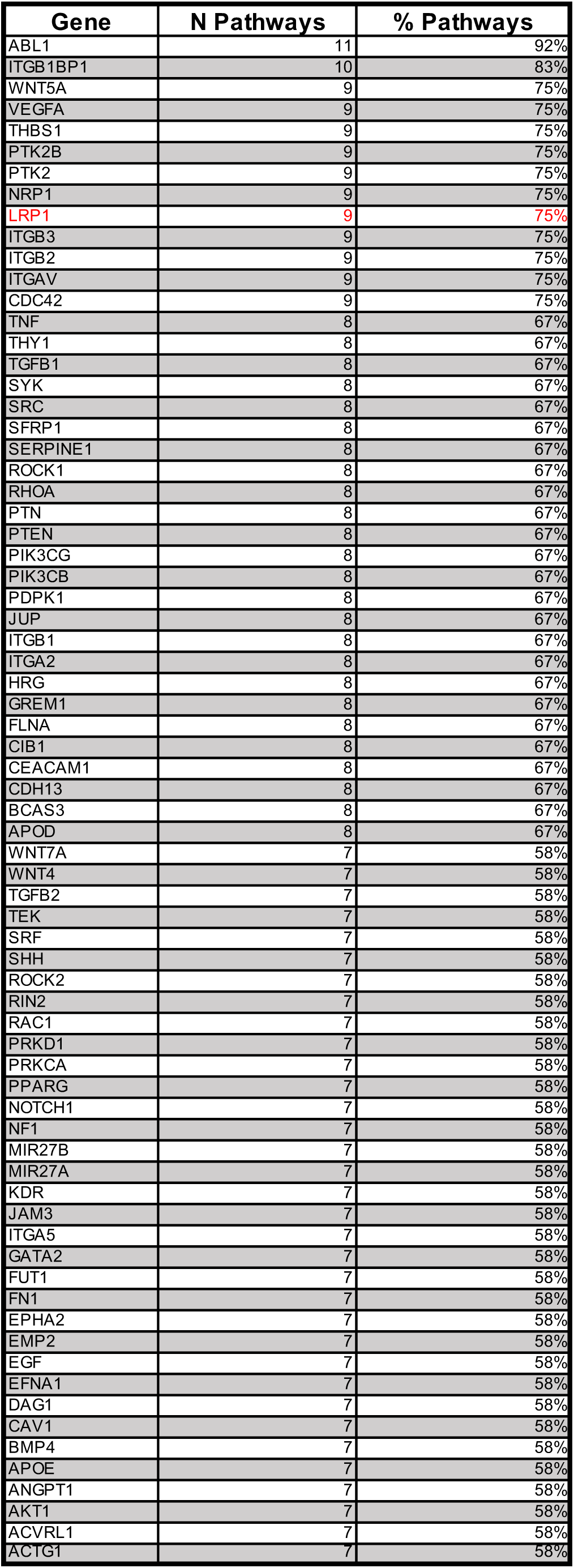

| Gene | N Pathways | % Pathways |
| --- | --- | --- |
| ABL1 | 11 | 92% |
| ITGB1BP1 | 10 | 83% |
| WNT5A | 9 | 75% |
| VEGFA | 9 | 75% |
| THBS1 | 9 | 75% |
| PTK2B | 9 | 75% |
| PTK2 | 9 | 75% |
| NRP1 | 9 | 75% |
| LRP1 | 9 | 75% |
| ITGB3 | 9 | 75% |
| ITGB2 | 9 | 75% |
| ITGAV | 9 | 75% |
| CDC42 | 9 | 75% |
| TNF | 8 | 67% |
| THY1 | 8 | 67% |
| TGFB1 | 8 | 67% |
| SYK | 8 | 67% |
| SRC | 8 | 67% |
| SFRP1 | 8 | 67% |
| SERPINE1 | 8 | 67% |
| ROCK1 | 8 | 67% |
| RHOA | 8 | 67% |
| PTN | 8 | 67% |
| PTEN | 8 | 67% |
| PIK3CG | 8 | 67% |
| PIK3CB | 8 | 67% |
| PDPK1 | 8 | 67% |
| JUP | 8 | 67% |
| ITGB1 | 8 | 67% |
| ITGA2 | 8 | 67% |
| HRG | 8 | 67% |
| GREM1 | 8 | 67% |
| FLNA | 8 | 67% |
| CIB1 | 8 | 67% |
| CEACAM1 | 8 | 67% |
| CDH13 | 8 | 67% |
| BCAS3 | 8 | 67% |
| APOD | 8 | 67% |
| WNT7A | 7 | 58% |
| WNT4 | 7 | 58% |
| TGFB2 | 7 | 58% |
| TEK | 7 | 58% |
| SRF | 7 | 58% |
| SHH | 7 | 58% |
| ROCK2 | 7 | 58% |
| RIN2 | 7 | 58% |
| RAC1 | 7 | 58% |
| PRKD1 | 7 | 58% |
| PRKCA | 7 | 58% |
| PPARG | 7 | 58% |
| NOTCH1 | 7 | 58% |
| NF1 | 7 | 58% |
| MIR27B | 7 | 58% |
| MIR27A | 7 | 58% |
| KDR | 7 | 58% |
| JAM3 | 7 | 58% |
| ITGA5 | 7 | 58% |
| GATA2 | 7 | 58% |
| FUT1 | 7 | 58% |
| FN1 | 7 | 58% |
| EPHA2 | 7 | 58% |
| EMP2 | 7 | 58% |
| EGF | 7 | 58% |
| EFNA1 | 7 | 58% |
| DAG1 | 7 | 58% |
| CAV1 | 7 | 58% |
| BMP4 | 7 | 58% |
| APOE | 7 | 58% |
| ANGPT1 | 7 | 58% |
| AKT1 | 7 | 58% |
| ACVRL1 | 7 | 58% |
| ACTG1 | 7 | 58% |

To address this hypothesis, a myc-tagged Gpr126 protein was stably overexpressed in iBECs and analyzed for mRNA levels of *Gpr126* and *Lrp1*. Overexpression of Gpr126 (by 6-fold) increased *Lrp1* mRNA levels (Supplementary Figure 6G). Then, using immunoprecipitation assays, we showed that in iBECs, Gpr126-myc co-immunoprecipitated both Lrp1 and β1-integrin (Figure 6A), and that a β1-integrin specific antibody co-immunoprecipitated with Gpr126-myc, which demonstrated the presence of Gpr126 and Lrp1 in the complex (Figure 6B). Furthermore, at the electron microscopy level, β1-integrin and Gpr126 were detected on the plasma membrane and in late endosomes of ECs in the brain cortex, which suggested that they are co-internalized and undergo trafficking to late endosomes and recycling to the plasma membrane, to promote cell migration (Figure 6C).

**Figure 6.**
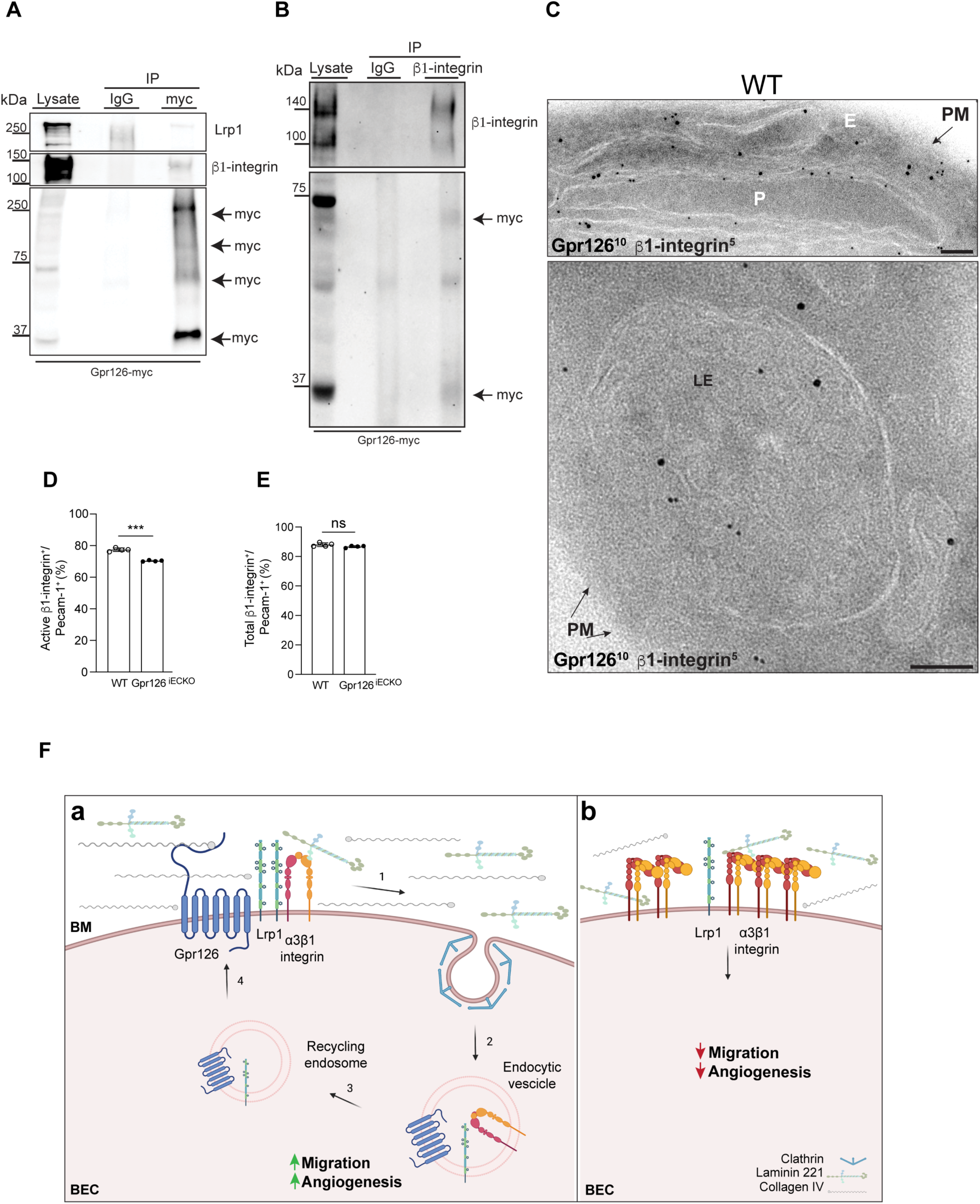
Gpr126 regulates angiogenesis by forming a complex with Lrp1 and α3β1-integrin. **A,** Representative immunoblotting for Lrp1, β1-integrin, and myc in immortalized brain endothelial cells (iBECs) transfected with Gpr126-myc. The protein extracts (lysates) were immunoprecipitated with an anti-myc antibody [IPs (myc)] or IgG [IPs (IgG)], as control. Data are representative of 3 independent experiments. **B,** Representative immunoblotting for β1-integrin and myc in iBECs transfected with Gpr126-myc. The protein extracts (lysates) were immunoprecipitated with an anti-β1-integrin antibody [IPs (β1-integrin)] or IgG as control. Data are representative of 3 independent experiments. **C,** EM of Gpr126 immunogold-labeled (10 nm gold, Gpr126^10^) and β1-integrin (5 nm gold, β1-integrin^5^) as representative cryo-sections of brain capillaries from WT mouse cortex at P18. PM, plasma membrane; E, endothelial cell; P, pericyte; LE, late endosome. Scale bars: 100 nm. **D, E,** Flow cytometric analysis of active and total β1-integrin-positive fBECs from WT and Gpr126^iECKO^ mice at P18. **D**, Active β1-integrin−/Pecam-1-positive cells ratio expressed as percentages. Each symbol represents a single experiment (n=3 WT and 3 Gpr126^iECKO^ mice for each independent experiment, as means ±SD). ***, *P*<0.0005 (unpaired t-tests with Welch’s correction). **E,** Total β1-integrin−/Pecam-1-positive cells ratio expressed as percentages. Each symbol represents a single experiment (n=3 WT and 3 Gpr126^iECKO^ mice for each independent experiment, as means ±SD). ns=not significant (unpaired t-tests with Welch’s correction). **F,** Schematic model for Gpr126 complex formation during angiogenesis. **(a)** Upon activation by collagen IV, Gpr126 binds to Lrp1 and α3β1-integrin (1). The complex goes through Lrp1-mediated endocytosis (2), which favours EC migration and angiogenesis. Then Lrp1 and Gpr126 are transported to the plasma membrane (3), through the recycling endosomes (4). **(b)** The absence of Gpr126 negatively affects BM deposition and expression of Lrp1. α3β1-integrin is not internalized, and hence it accumulates on the surface. These processes result in a reduction of migration and angiogenesis. BEC, brain endothelial cell; BM, basement membrane.

Moreover, based on the activity status, it has been reported that integrins undergo trafficking with significantly different kinetics, which has been attributed to changes in cell migration and invasion (Arjonen et al., 2012). As a consequence, the active form of β1-integrin was predominantly localized in endosomes during migration, while the inactive integrin was in the detached membrane protrusions on the plasma membrane, where it might be involved in adhesion processes.

The absence of Gpr126 impaired EC migration (Figure 4D-F and Supplementary Figure 4J, K), which might be caused by reduced β1-integrin activation. To explore this hypothesis, we performed fluorescence-activated cell sorting, as a comparison of fBECs of WT and *Gpr126^iECKO^* pups. Although the total β1-integrin was similar, the active form was reduced in the absence of Gpr126 (Figure 6D and E). As it is well established that LRP1-mediated endocytosis of β1-integrin (Rabiej et al., 2016) and the expression of Lrp1 is reduced in Gpr126*^iECKO^*cells (Figure 5D-F), multiple lines of evidence converged to support the involvement of the Gpr126-Lrp1 complex in cell migration, to balance the levels of adhesiveness through β1-integrin activation and recycling. To identify which α-subunit coupled to β1-integrin, we showed that Gpr126-myc co-immunoprecipitated with α3β1-integrin, but not with the α1 subunit (Supplementary Figure 6H).

Altogether, our findings indicate the involvement of the Gpr126–Lrp1–α3β1-integrin complex in EC migration during angiogenesis, as summarized in Figure 6F.

### Gpr126 deficiency in the adult is protective after acute damage to the CNS

So far, the present study has demonstrated that Gpr126 expression is required for correct postnatal development of the BBB. Therefore, we investigated whether the impairment of BBB properties induced by Gpr126 inactivation at P1-P4 was still present in the adult mouse brain vasculature (i.e., at 2 months old). Adult *Gpr126^iECKO^* mice with 75% Gpr126 depletion (Supplementary Figure 7A, B), were viable and appeared healthy for up to 1 year. Indeed, these mice did not show any major defects in vessel morphology or permeability, as measured by the tracer cadaverine extravasation in the brain, which indicated that the vascular barrier was correctly formed (Supplementary Figure 7C-E). Under electron microscopy, no significant changes in the BM distribution were detected in the brain cortex between the *Gpr126^iECKO^* mice and their WT littermates (Supplementary Figure 7F). Of note, in the retinal vasculature, there was persistence of frequent arteriovenous crossovers in the absence of Gpr126 (Supplementary Figure 7G, H). These data suggested that under physiological conditions a compensatory mechanism develops in response to the absence of Gpr126 over time in the adult brain.

We next asked whether inactivation of Gpr126 at P1-P4 still impact acute CNS pathologies where the BBB is strongly damaged. We thus induced transient ischemic stroke using the middle cerebral artery occlusion (MCAo) in the adult brain (Figure 7A).

**Figure 7.**
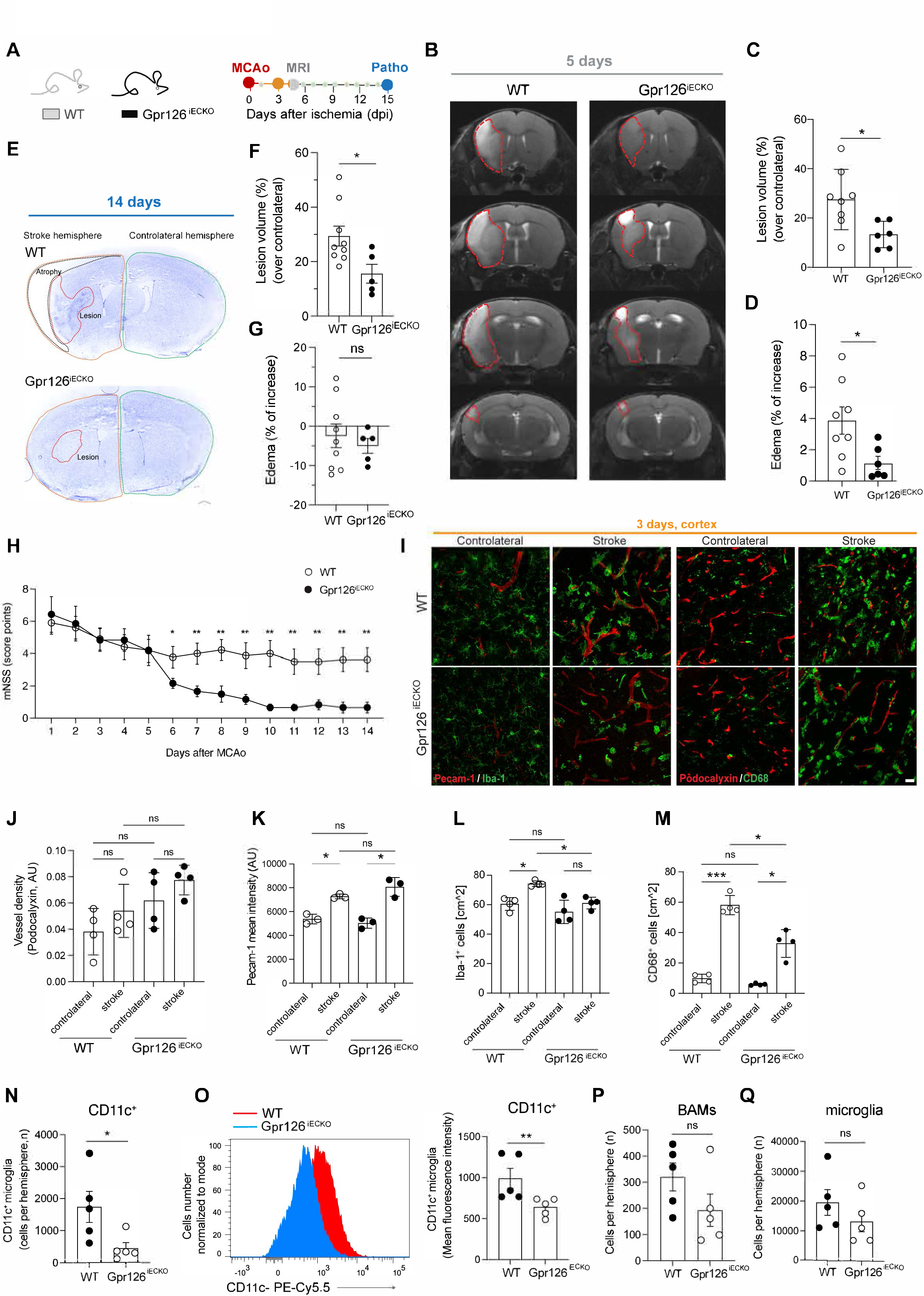
Gpr126 deficiency is protective after acute damage to the CNS. **A,** Schematic experimental design of the stroke study in adult WT and Gpr126^iECKO^ mice (2 months old). **B,** Magnetic resonance imaging of mouse brains from WT and Gpr126^iECKO^ mice 5 days after MCAo. Red dashed line indicates the stroke area. **C,** Quantification of lesion volume (white, left hemisphere) corrected for edema and expressed as a percentage of the contralateral non-lesioned area (right hemisphere), as shown in **B**. Each symbol represents a mouse (n=8 WT and 6 Gpr126^iECKO^ mice [1 Gpr126^iECKO^ mouse was excluded from the analysis as an outlier; Grubb’s method, alpha = 0.2], as means ±SD). *, *P*≤0.05 (unpaired t-tests with Welch’s correction). **D,** Quantification of edema, expressed as a percentage of the contralateral nonlesioned area, as shown in **B**. Each symbol represents a mouse (n=8 WT and 6 Gpr126^iECKO^ mice, as means ±SD). *, *P* ≤0.05 (unpaired t-tests with Welch’s correction). **E**, Representative coronal sections of mouse brain from WT and Gpr126^iECKO^ mice stained with cresyl violet 14 days after MCAo. Infarct area is white (unstained by cresyl violet); orange and green dashed lines show the ipsilateral and contralateral hemispheres, respectively. Black line, brain atrophy in the WT mouse brain; red dashed line, lesion size. **F,** Quantification of lesion volume corrected for edema and expressed as percentage of the contralateral nonlesioned area, as shown in **E**. Each symbol represents a mouse (n=9 WT and 5 Gpr126^iECKO^ mice, as means ±SD). *, *P* ≤0.05 (unpaired t-tests with Welch’s correction). **G,** Quantification of edema expressed as percentage of the contralateral nonlesioned area, as shown in **E**. Each symbol represents a mouse (n=9 WT and 5 Gpr126^iECKO^ mice, as means ±SD). ns=not significant (unpaired t-tests with Welch’s correction). **H,** Clinical disability measured by modified neurological stroke scale (mNSS) in WT and Gpr126^iECKO^ mice 14 days after MCAo (n=10 WT and 7 Gpr126^iECKO^ mice, as means ±SD). *, P ≤0.05; **, P ≤0.005 (uncorrected Fisher’s LSD multiple comparisons). **I,** Representative confocal images of brain cortex cryosections (10 μm, contralateral and ipsilateral) from WT and Gpr126^iECKO^ mice 3 days after MCAo. Podocalyxin (red) and Pecam-1 (red) stained the vasculature. Iba-1 (green) and CD68 (green) stained the microglia. Scale bar: 10 μm. **J,** Quantification of vessel density in brain cortex cryosections, as shown in **I**. Each symbol represents a mean of 6 sections for each mouse (n=4 WT and 4 Gpr126^iECKO^ mice, as means ±SD). ns=not significant, Brown-Forsythe and Welch ANOVA, Dunnett’s T3 multiple comparisons tests). **K,** Quantification of Pecam-1 signal (mean intensity) in the vessels of brain cortex cryosections (10 μm) as shown in **I**, expressed as arbitrary units (AU). Each symbol represents a mean of 6 sections for each mouse. (n=3 WT and 3 Gpr126^iECKO^ mice, as means ±SD). ns=not significant; *, P ≤0.05 Brown-Forsythe and Welch ANOVA, Dunnett’s T3 multiple comparison tests). **L,** Quantification of the number of cells expressing Iba-1 per area, as shown in **I**. Each symbol represents a mean of 6 sections for each mouse (n=4 WT and 4 Gpr126^iECKO^ mice, as means ±SD). ns=not significant; *, P ≤0.05 (Brown-Forsythe and Welch ANOVA, Dunnett’s T3 multiple comparisons tests). **M,** Quantification of the number of cells expressing CD68 per area, as shown in **I**. Each symbol represents a mean of 6 sections for each mouse (n=4 WT and 4 Gpr126^iECKO^ mice, as means ±SD). ns=not significant, *, P ≤0.05, ***, *P*<0.0005 (Brown-Forsythe and Welch ANOVA, Dunnett’s T3 multiple comparison tests). **N-Q,** Flow cytometry analysis of microglia cells in the stroke hemisphere from WT and Gpr126^iECKO^ mice 3 days after MCAo. **N**, Quantification of the number of CD11c-positive cells. Each symbol represents a mouse (n=5 WT and 5 Gpr126^iECKO^ mice for each independent experiment, as means ±SD). *, P ≤0.05 (unpaired t-tests with Welch’s correction). **O,** Representative plot (left) and quantification of CD11 signal on microglia cells (right) expressed as mean fluorescence intensity. Each symbol represents a mouse (n=5 WT and 5 Gpr126^iECKO^ mice for each independent experiment, as means ±SD). **, P ≤0.005 (unpaired t-tests with Welch’s correction). **P**, **Q,** Quantification of border-associated macrophages (BAMs) (**P**) and microglia (**Q**) cells. Each symbol represents a mouse (n=5 WT and 5 Gpr126^iECKO^ mice for each independent experiment, as means ±SD). ns=not significant (unpaired t-tests with Welch’s correction).

Examination of a publicly available dataset of a single-cell transcriptome dataset (Zheng et al., 2022) showed that in healthy mouse adult brain, *Gpr126* is mainly expressed in veins, rather than arteries and capillaries (Supplementary Figure 8A). No changes in *Gpr126* expression were seen after MCAo, compared to the sham counterpart (Supplementary Figure 8B).

Interestingly, at 5 days after MCAo, *Gpr126^iECKO^* mice showed significantly reduced ischemic lesion volumes and decreased edema, compared to the WT mice (Figure 7B-D). These diminished ischemic lesion volumes were confirmed by histology at 14 days after ischemia, as the end of the follow-up (Figure 7E-G). At this later time, the early edema was resolved and brain tissue atrophy ensued. Accordingly, *Gpr126^iECKO^* mice had reduced behavioral disability compared to the WT mice, as assessed by the modified neurological severity scores (Figure 7H). These data demonstrated a rather protective effect of Gpr126 deficiency on lesion burden and neurological dysfunction in acute ischemic stroke conditions.

Using histology and single cell cytofluorimetry, we then investigated the effects of endothelial Gpr126 absence in ischemic stroke on the brains at 3 days post-MCAo. The vascular density and morphology were similar in the stroke area of *Gpr126^iECKO^* mice, compared to WT (Figure 7I, J), while Pecam-1 expression was significantly increased upon stroke in both WT and *Gpr126^iECKO^* mice, compared to their contralateral cortices; the absence of Gpr126 did not prevent this increase in Pecam-1 expression (Figure 7I, K).

We then explored inflammatory cells in the contralateral cortex and in the stroke area, through analysis of microglia Iba1^+^ and CD68^+^ cells by histology. The absence of Gpr126 in the contralateral hemisphere did not change the numbers of Iba-1^+^ and CD68^+^ cells, compared to WT, nor typical highly ramified morphology of Iba-1^+^ cells, which showed a small soma and fine and long processes (Figure 7I, L). In contrast, in the stroke cortex area of Gpr126^iECKO^ mice, the numbers of Iba-1^+^ and CD68^+^ cells were significantly reduced, compared to WT, while the change in Iba-1^+^ cell morphology from highly ramified to ameboid was comparable (Figure 7I-M).

To investigate in more detail the inflammatory cells after stroke, we also performed 18-color single cell cytofluorimetric analysis. No significant differences were seen for B-cells, CD4^+^ and CD8^+^ T cells, dendritic cells, neutrophils, and monocytes (Supplementary Figure 8C-H). However, the numbers of activated CD11c^+^ microglia cells were significantly reduced in *Gpr126^iECKO^* mice, also in terms of the CD11c expression levels (as mean fluorescence intensity), compared to WT (Figure 7N, O). Furthermore, in *Gpr126^iECKO^* mice, there was a decrease in border associated macrophages, although this effect did not reach significance (Figure 7P, Q). These data suggested a detrimental role of EC Gpr126 expression in the activation of microglia following MCAo, which will contribute to the bad outcomes seen after brain ischemic stroke.

Taken together, these data show that lack of Gpr126 attenuates microglia activation upon MCAo, to contribute to improved neurological outcome after ischemic stroke.

## Discussion

Central nervous system development, angiogenesis and BBB maturation require regulation of multiple pathways activated by distinct signals. These multi-step processes appear to initiate with the neural-derived Wnt induction of BBB properties during the angiogenic program, in combination with pericytes, astrocytes and environmental cues. While individual cellular and molecular interactions have been identified, how the downstream effectors of Wnt signaling induce angiogenesis during development to regulate different features of the BBB needs further investigation.

Adhesion GPCRs form a large family of GPCRs that are essential for multiple biological processes. Lack of knowledge about the functions of specific receptors has hampered their development as therapeutic drug targets (Yang et al., 2021). Here, we reveal a previously unknown role for the adhesion receptor Gpr126 in the brain microvasculature. We have demonstrated that Gpr126 is a novel target of Wnt/β-catenin signaling and is required for postnatal BBB development; surprisingly, Gpr126 expression is also detrimental to ischemic stroke in the adult.

In recent years, the expression of Gpr126 in ECs has been under debate. Several reports have shown that Gpr126 is expressed in human umbilical vein and aortic ECs (Patra et al., 2013; Stehlik et al., 2004) and in the endothelium in a variety of organs (e.g., lung, tongue, kidney, stomach, endocardium) and at different developmental stages (e.g., embryo, P5, adult; using a reporter mouse model (Musa et al., 2019). However, these studies failed to detect Gpr126 in the brain microvasculature.

Here, we have shown that murine Gpr126 expression levels increase progressively in early postnatal life, to reach a peak of expression at P18, when establishment of the BBB is reported (Biswas et al., 2020). Moreover, as previously published, Gpr126 expression is low in the embryo, at early postnatal stages (P2-P11), and in the adult (Musa et al., 2019; Waller-Evans et al., 2010).

An important finding from the present study is that Gpr126 is one of the main players downstream of Wnt/β-catenin signaling during development, in the control of cerebral angiogenesis and BBB genesis. Postnatally induced Gpr126 inactivation led to vasculature alterations in the brain cortex and retina, and induced a sparser network of enlarged and leaky vessels. This vessel enlargement is not linked to EC hyperplasia, but to a decrease in the thickness of the ECs forming the vessel. The vessel swelling that occurs in the absence of Gpr126 might also be related to the loss of the vascular smooth muscle cells and the pericytes around them, as these cells regulate the vascular tone (Mae et al., 2021). Moreover, several studies have demonstrated that the deposition of BM proteins also leads to restrictions in vessel diameter and patterning, while disruption of BM assembly and EC–pericyte interactions leads to unregulated and mispatterned vessel growth, including increased vessel width (Stratman et al., 2009; Stratman et al., 2010). In addition, there is evidence that Gpr126 acts through cell– cell and cell–ECM interactions upon binding to collagen IV or laminin-211 (Paavola et al., 2014; Petersen et al., 2015).

Here, we identified the adhesion receptor Gpr126 as a key signaling molecule that is required for EC–pericyte interactions, to correctly assemble the BM matrix proteins (i.e., collagen-IV, fibronectin, a2 laminin), to ensure correct vascular morphogenesis and maturation during blood-barrier development in the brain and in the retina. Moreover, the conspicuous structural abnormalities of the BM, which include a loose association with ECs and pericytes, can contribute to vessel regression (Baluk et al., 2003). Vessel ghosts indicate a regressing segment, and these are found at the advancing front of an angiogenic network in the retina of *Gpr126^iECKO^* mice at P18, along with regression events that can be appreciated in *in-vitro* angiogenesis assays. This analysis suggests that Gpr126 has a role in stabilizing newly formed vessels, to guide them into the spatially distributed and functional network required for BBB development.

An earlier study reported a requirement for GPR126 in angiogenesis, through regulation of the signaling pathway of VEGFR2 in human dermal microvascular ECs (Cui et al., 2014). However, our data confirm the involvement of Gpr126 in angiogenesis and BBB development, through down-regulation of *Vegfa* expression (as a ligand of Vegfr2) and other key genes involved in angiogenesis processes in the brain vasculature of *Gpr126^iECKO^* mice, compared to WT. Among these genes, our transcriptomic data also reveal down-regulation of *Lrp-1* mRNA in the absence of Gpr126.

Interestingly, recent studies have linked LRP1 with EC growth, migration, and angiogenesis, although the molecular mechanisms remain largely unknown (Nakajima et al., 2014; Pi et al., 2012). LRP1 loss-of-function animal models, such as zebrafish with morpholino knockdown and tissue-specific knockout mice, showed vascular developmental defects with an interrupted endothelial layer and extensive hemorrhage (Nakajima et al., 2014; Pi et al., 2012). There is also evidence that mouse embryonic fibroblasts and neuronal cells with a knock-in mutation of the LRP1 NPxY motif (that is responsible for β1-integrin interactions) show impaired cell migration (Avraamides et al., 2008; Rabiej et al., 2016). Our data demonstrate that upon collagen IV binding, Gpr126 sustains the expression of Lrp1 through a novel regulatory loop, and for the first time we show that Gpr126 is required during developmental angiogenesis, to promote migration via Lrp1 and α3β1-integrin interactions.

Overall, these data highlight a role for Gpr126 as an important regulator of angiogenesis and barrier genesis during BBB development.

In contrast, in the adult brain, vascular morphogenesis and permeability did not show any major changes when comparing Gpr126 deficient and control mice, while we observed frequent arteriovenous crossovers in the retina of *Gpr126^iECKO^* mice. The mechanisms remain undetermined. Arteriovenous crossovers have also been reported in β-catenin, VEGF-A (Haigh et al., 2003), Nrp1 (Fantin et al., 2011), and VEGFR2 (Zarkada et al., 2015) loss-of-function models. The persistence of vascular misspatterning in the retina might depend on the timing of Gpr126 deletion during BBB development. Indeed, *Gpr126* gene inactivation was induced postnatally, while angiogenesis and barrier genesis in the brain have already started in the embryo, while in the retina they occur entirely after birth. However, we still do not know the factors that induce the mechanism that compensates the phenotype in the brain adult mice under physiological conditions.

Recently, a mechanism termed transcriptional adaptation was described in mice, by which aberrant mRNA degradation triggers up-regulation of a potential compensatory gene or genes that can provide normal function (El-Brolosy et al., 2019). Thus, to identify potential compensatory paralogs or “adapting genes” that attenuate the BBB defects in adult *Gpr126* mutants, we previously analyzed the expression of genes with high sequence similarities to mouse *Gpr126*, including *Gpr64* and *Gpr133* (Patra et al., 2014). Interestingly, BECs did not express both *Gpr64* and *Gpr133* and the lack of *Gpr126* did not promote up-regulation of these potential compensatory genes. Thus, our understanding of the underlying molecular mechanisms that control this process still remains limited.

Also recently, in a multi-trait analysis, the *GPR126* locus was associated with lacunar stroke with genome-wide significance, although any loss or gain of function of the gene has been reported (Traylor et al., 2021). Here, we report that the lack of endothelial Gpr126 in adult mice is protective in acute ischemic stroke, which leads to a reduced brain lesions and edema, and a decreased neurological disability. Therefore, these data underline that Gpr126 is not required to preserve the integrity of the brain vasculature under pathological ischemic conditions, but rather contributes to injury exacerbation. Moreover, in 2016, Mogha et al. showed that Iba-1^+^ and CD68^+^ cell recruitment was impaired in a conditional knockout Gpr126 mouse, following nerve sciatic crush (Mogha et al., 2016). The brain resident macrophages, the microglia, were activated several minutes after ischemic stroke, and this activation peaked after 2–3 days (Jin et al., 2013). Activated microglia have deleterious effects on the acute phase of ischemic stroke due to their release of cytotoxic factors, such as reactive oxygen species, tumor necrosis factor-alpha, and interleukin 1-beta, which leads to BBB disruption and activation of post-stroke inflammation. Here, we found that the absence of Gpr126 in BECs specifically attenuated microglial activation and CD68^+^ cell recruitment in the ischemic lesion, without affecting the overall number of immune cells at the sites of injury (including neutrophils, monocytes, and lymphocytes). Furthermore, analysis of vessel morphology and density did not show major changes when comparing Gpr126 deficient and control mice 3 days post-stroke. One possible explanation is that in the absence of Gpr126, the brain vasculature inhibits the release of pro-inflammatory molecules that prevent microglial activation, thus improving the outcome of a stroke.

Therefore, we propose Gpr126 as a promising therapeutic target to be considered in ischemic stroke. Progress in the understanding of the signaling mechanisms involved in the therapeutic effects of Gpr126 inhibition might contribute to the development of novel drugs for the treatment of patients with ischemic stroke.

## Materials and Methods

### Mice

The following mouse strains were used: Gpr126^flox/flox^ (model TF0269; Taconic Biosciences GmbH), *Cdh5(PAC)-CreER^T2^*(13073; Taconic Biosciences GmbH), Rosa26-Stopflox-EYFP (provided by Stefano Casola, IFOM ETS - The AIRC Institute of Molecular Oncology), C57BL/6J mice (Charles River, Italy). Gpr126^flox/flox^ mice were generated by Lexicon Pharmaceuticals (The Woodlands, TX, USA). Exons 3 and 4 of the *Gpr126* gene were floxed with LoxP sequences. A neomycin (NEO) cassette flanked by FRT sites was included in the floxed *Gpr126* sequence. Heterozygous animals were obtained on a mixed background (129/SvEv-C57BL/6). The marker-assisted accelerated backcrossing method (MAX-BAX; Charles River) was used to obtain Gpr126^flox/flox^ strain on a pure C57BL/6J background.

The Gpr126^flox/flox^ mice were crossed with *Cdh5(PAC)-CreER^T2^*mice to generate a *Gpr126* conditional knock-out (Gpr126^iECKO^), and then crossed with Rosa26-Stopflox-EYFP mice to detect recombination of the floxed allele.

Tamoxifen (T5648; Sigma-Aldrich) was dissolved in 10% ethanol-sunflower oil (10 mg/mL) and administrated to the mice to induce Cre activity and genetic modifications. For inducible endothelial-specific deletion of Gpr126, *Cdh5(PAC)-CreER^T2^-*Gpr126^flox/flox^ mouse models were injected subcutaneously with tamoxifen at the age of post-natal day (P) 0, P1, P2, P3. All of the mice were genotyped to verify the mutations, as previously reported [for Gpr126 (Mogha et al., 2013); for Cre (Liebner et al., 2008)]. All of the animal procedures were in accordance with the Institutional Animal Care and Use Committee, and in compliance with the guidelines established in the Principles of Laboratory Animal Care (directive 86/609/EEC); they were also approved by the Italian Ministry of Health.

### Cells and treatments

#### Freshly isolated brain endothelial cells (fBECs) and primary cultured (c)BECs

WT and Gpr126^iECKO^ ECs were derived from brains of embryos and pups at different post-natal stages upon tamoxifen treatment, as previously reported. Briefly, the brains were digested enzymatically in combination with gentle dissociation (MACS; Neural tissue dissociation kit [130-092-628] for embryos and pups ≤P7; adult brain dissociation kit [130-107-677] for pups P8 and older; Miltenyi Biotech). After dissociation, the myelin cell debris and erythrocytes were removed according to the manufacturer protocol. ECs were enriched by depletion of CD45-positive cells with CD45 microbeads (30-052-301; Miltenyi Biotech). After AN2 re-expression according to the manufacturer protocol, pericytes and astrocytes were depleted using AN2 microbeads (130-097-171; Miltenyi Biotech) and ACSA-2 microbeads (130-097-678; Miltenyi Biotech), respectively, followed by positive selection using CD31 microbeads (30-097-418; Miltenyi Biotech).

Alternatively, EC isolation and cultured brain microvascular fragments (referred to as primary cBECs in the main text) were processed as previously described (Calabria et al., 2006; Liebner et al., 2000). Capillary fragments were seeded into collagen I (354236; BD Biosciences)-coated wells (0.5 brains/4 cm^2^) and cultured in Dulbecco’s modified Eagle’s medium (DMEM) plus GlutaMAX (10564011; ThermoFisher Scientific), 20% fetal bovine serum (Hyclone), 100 mg/mL heparin (H3149; Sigma-Aldrich), and 5 mg/mL Endothelial Cell Growth Supplement (E2759; Sigma-Aldrich). After 3 days of puromycin selection (4 mg/mL; AG-CN2-0078; Adipogen), cBECs were treated with purified Wnt3a (100 ng/mL; 1324-WN; R&D) or purified Wnt5a (250 ng/mL; 645-WN; R&D), in complete medium, every other day for 5 days. Alternatively, cBECs cells were treated with L-supernatant or Wnt3a-conditioned medium diluted 1:2 in complete medium (Paolinelli et al., 2013).

For Gpr126 receptor activation, WT and Gpr126^iECKO^ cBECs were treated with 5 μg/mL collagen-IV (3410-010-01; Cultrex) or phosphate-buffered saline (PBS) for 2 days in DMEM plus GlutaMAX and 1% fetal bovine serum (Hyclone, starving medium). cBECs were cultured in starving medium for 16 h prior to treatment for 45 min with collagen-IV. The final cell pellets were washed with PBS and processed for protein or RNA extraction.

#### Lung endothelial cells in culture (cLEC)

ECs were derived from lungs of WT adult mice (8 weeks old). Briefly, the lungs were digested enzymatically in combination with gentle dissociation (MACS; lung dissociation kits; 130-095-927; Miltenyi Biotech). After dissociation, the ECs were enriched by depletion of CD45-positive cells with CD45 microbeads (30-052-301; Miltenyi Biotech). After AN2 and CD326 re-expression according to the manufacturer protocol, immune cells, lymphatic ECs, fibroblasts, epithelial cells, smooth muscle cells, and pericytes were removed using the following microbeads: anti-CD45 (130-052-301), anti-CD90.2 (130-121-278), anti-CD326 (130-105-958), anti-CD138 (130-098-257) and anti-AN2 (130-097-170). Finally, the ECs were positively selected using CD31 microbeads (30-097-418; Miltenyi Biotech). LECs were seeded into 0.1% gelatin (214340; Difco)-coated wells (1 lung/2 cm^2^) and cultured in DMEM plus GlutaMAX (10564011; ThermoFisher Scientific), 20% fetal bovine serum (Hyclone), 100 mg/mL heparin (H3149; Sigma-Aldrich), and 5 mg/mL Endothelial Cell Growth Supplement (E2759; Sigma-Aldrich). After 5 days, cLECs were treated with purified Wnt3a (100 ng/mL; 1324-WN; R&D), in complete medium described above, every other day for 5 days. The final cell pellets were washed with PBS and processed for protein or RNA extraction.

#### Immortalized brain endothelial cells (iBECs)

Brain ECs were isolated from Gpr126^flox/flox^ pups (P9) (as described above for cBECs), and then seeded into collagen I (354236; BD Biosciences)-coated wells (1 brain/2 cm^2^) and cultured in MCDB-131 (10372-019; ThermoFisher Scientific), 20% fetal bovine serum (Hyclone), 100 mg/mL heparin (H3149; Sigma-Aldrich), and 5 mg/mL Endothelial Cell Growth Supplement (E2759; Sigma-Aldrich) (complete medium). After a day, the cells were immortalized in culture through retroviral expression of polyoma middle T antigen, as described previously (Balconi et al., 2000), and then left to grow for 4-5 weeks, changing the medium every 3-4 days. Gpr126 was inactivated by treating the cultured iBECs with TAT-Cre recombinase (100 μg/mL) and chloroquine (50 μM) (C6628; Sigma-Aldrich) for 1 h in Hyclone medium. Finally, the cells were cultured in complete medium.

### Cadaverine permeability assay

Lysine-fixable, Alexa Fluor-555 conjugated cadaverine (3.125 mg/mL in saline; 950 Da; A30677; ThermoFisher Scientific) was injected intraperitoneally in mice at P18 and adults (25 mg/kg). The circulation time was 2 h. The animals were anesthesised by intraperitoneal injection of avertin (20 mg/kg; T48402; Sigma-Aldrich) 20 min before sacrifice. Then, the mice were perfused for 1 min to 2 min with Hank’s balanced salt solution (14025-050; ThermoFisher Scientific), followed by 5 min perfusion with 4% paraformaldehyde (PFA) in PBS, pH 7.2. The brains from 3 animals per group were dissected out and fixed for up to 8 h by immersion in 4% PFA at 4 °C, then washed in PBS, and prepared for vibratome sectioning. Brains were fully sectioned. Sagittal vibratome sections were incubated with anti CD93 (AF1696, R&D Systems) in blocking/permeabilisation solution (5% donkey serum [017-000-1210; Jackson Immuno Research], 0.3% Triton X-100, in PBS) overnight at 4 °C. This was followed by incubation with anti-sheep Alexa Fluor 488 (713-545-147; Jackson Immuno Research). For more details see Tissue processing and immunohistochemistry.

### Lentiviral preparation

To stably express Gpr126, a custom-made lentiviral vector was purchased from Vector Builder. This is a *bicistronic* expression *vector* that uses a self-cleaving *2A peptide* to co-express murine Adgrg6 and EGFP under the control *of* a single CMV promoter. The lentiviruses were generated in HEK 293T cells, as described previously (Morini et al., 2018). The lentiviral vectors were produced (Dull et al., 1998) with the packaging plasmids donated by L. Naldini (HSR-TIGET, San Raffaele Telethon Institute for Gene Therapy, Milan, Italy). Infectious viruses were purified and titred using standard techniques. Two consecutive cycles of infections were performed in complete medium.

### *In-situ* hybridization using the RNAscope technique

*In-situ* hybridization using the RNAscope technique was performed according to the manufacturer instructions (Advanced Cell Diagnostics), using multiplex fluorescent Reagent Kit v2. Briefly, brains derived from WT and Gpr126^iECKO^ mice were dissected out and post-fixed in 4% formaldehyde overnight at 4 °C, and then processed using the standard paraffin-embedding procedure.

For paraffin-embedding, the brains were processed using a Diapath automatic processor, as follows. Tissues were dehydrated through 70% (60 min), 2 changes of 95% (90 min each), and 3 changes of 99% (60 min each) ethanol, cleared through 3 changes of xylene (90 min each), and finally immersed in 3 changes of paraffin (1 h each). Samples were embedded in a paraffin block and prepared for sectioning. Four-μm-thick sagittal sections were deparaffinized and pretreated with H_2_O_2_, followed by target retrieval and a 15 min incubation with protease IV. The probes *Cldn5* (#491611-C2; Advanced Cell Diagnostics) and *Gpr126* (#1038161-C1; Advanced Cell Diagnostics) were hybridized on the tissue sections for 2 h at 4 °C (HybEZ oven; Advanced Cell Diagnostics), followed by signal amplification with the reagents included in Reagent Kit v2. The signal was detected for *Cldn5* with OPAL520 (SKU FP1487001KT; Akoya Bioscience) and for *Gpr126* with OPAL650 (SKU FP1496001KT; Akoya Bioscience), and tissue sections were counterstained with DAPI (Advanced Cell Diagnostics). Fluorescent tilescan images that visualized the signals for mRNA expression were acquired using a fluorescent microscope (DMi8; Leica), at 40× magnification. Representative close-up images were then acquired with the confocal microscope at 63× magnification. For comparison purposes, different sample images of the same probe combinations were acquired under constant acquisition settings.

### Gene expression analysis

#### Affymetrix

Total mRNA was extracted from cBECs after treatment with Wnt3a. Gene expression was analyzed using the GeneChip ST 1.0 array (Affymetrix) covering 29,000 murine genes. For each condition, mRNA obtained from three different experiments was assayed to account for biological variability. Genes differentially expressed between the two conditions were identified as those having at least a two-fold change and P<0.05 using Fisher’s least significant difference tests.

#### RNA sequencing

Total RNA was extracted from fBECs of WT and Gpr126^iECKO^ mice at P18 using Maxwell RSC simplyRNA tissue kits with the *Maxwell RSC* Instruments (Promega Corporation). The amount of RNA was measured using the Nanodrop technique and its integrity assessed using Agilent Bioanalyzer 2100 with Nano RNA kits (RIN > 8). Libraries for RNA sequencing were prepared following the manufacturer protocols for transcriptome sequencing with the Illumina NextSeq 550DX sequencer (Illumina). mRNA-seq indexed library preparation was performed starting from 500 ng of total RNA with the Illumina ligation stranded mRNA (Illumina) according to the manufacturer instructions. Indexed libraries were quality controlled on an Agilent Bioanalyzer 2100 with High Sensitivity DNA kits, quantified with Qubit HS DNA, normalized and pooled to perform multiplexed sequencing runs. A PhiX (1%) control was added to the sequencing pool, to serve as a positive run control. Sequencing was performed in PE mode (2×75nt) on an Illumina NextSeq550Dx platform, to generate on average 50 million PE reads per sample, on average.

#### RNA-sequencing data analysis

Reads were aligned to the mm10 assembly mouse reference genome using the STAR aligner [v 2.6.1d (Dobin et al., 2013)] and were quantified using Salmon [v1.4.0 (Patro et al., 2017)]. Differential gene expression analysis was performed using the Bioconductor package DESeq2 [v1.30.0 (Love et al., 2014)] that estimates variance-mean dependence in count data from high-throughput sequencing data and tests for differential expression exploiting a negative binomial distribution-based model.

Preranked gene set enrichment analysis (GSEA) for evaluating pathway enrichment in transcriptional data was carried out using the Bioconductor package fgsea (Sergushichev et al., 2016), taking advantage of the GO Biological process gene sets available from the GSEA Molecular Signatures Database (https://www.gsea-msigdb.org/gsea/msigdb/genesets.jsp?collections).

#### Quantitative RT-PCR analysis

Total RNA was extracted from fBECs from WT and Gpr126^iECKO^ mice at P18 using Maxwell RSC simplyRNA tissue kits with a *Maxwell RSC* instrument (Promega Corporation). The amount of RNA was measured using the Nanodrop technique and reverse transcribed with random hexamers (High-Capacity cDNA Archive kits; Applied Biosystems), following the manufacturer instructions. For gene expression analysis, 5 ng cDNA was amplified (in triplicate) in a reaction volume of 10 μL that contained the following reagents: 5 μL “TaqMan Fast Advanced Master Mix (4444557; Applied Biosystems), 0.5 μL TaqMan Gene expression assay 20× (ThermoFisher Scientific). For the time course of Gpr126 expression, the samples were preamplified using TaqMan PreAmp Master Mix (ThermoFisher Scientific), then diluited 1:4 with TE 1× buffer, and 2 μL/well was used for RT-PCR analysis. Real-time PCR was carried out on a real-time PCR system (7500; ThermoFisher Scientific) using a pre-PCR step of 20 s at 95 °C, followed by 4 cycles of 1 s at 95 °C and 20 s at 60 °C. A specific TaqMan assay (ThermoFisher Scientific) was used for each gene. Raw data (Ct) were analyzed using the Biogazelle qbase plus software, and the fold-changes are expressed as calibrated normalized relative quantities (CNRQs) with standard error (SE). The GeNorm Software chose GAPDH and 18s as the best housekeeping genes, and the geometric mean of GAPDH and 18s was used to normalise the data.

### Electron microscopy (EM)

WT and Gpr126^iECKO^ mice were anesthetized by intraperitoneal injection of avertin 20 mg/kg (T48402, Sigma) and perfused with 4% PFA and 2.5% glutaraldehyde (EMS, USA) mixture in 0.2 M sodium cacodylate pH 7.2 at the physiological pressure for 5 min (1.8 mL/min for adult mice and 0.6 mL/min for pups at p18). Brain cortex was excised and processed for EM examination and EM tomography as previously described (Beznoussenko et al., 2015). For immuno-labeling, the grids were prepared as previously described (Beznoussenko et al., 2022). Briefly, the grids were incubated for 10 min in 0.5% BSA-c in PBS (bovine serum albumin-c^TM^, acetylated, 10% in water; Aurion) to prevent nonspecific labeling, and then incubated on 10 μL droplets of primary anti-Gpr126 (ab218046; Abcam) diluted 1:10 for 2 h, and then with protein-A gold 10 nm (PAG10, CMC, Utrecht, The Netherlands) 1:50 diluted in blocking solution for 20 min at room temperature. The grids were rinsed six times with 0.1% BSA-c in PBS and post-fixed with 1% glutaraldehyde in 0.15 M HEPES (pH 7.3) for 5 min. The primary antibody was monoclonal anti-β1-integrin (AF 2325; R&D Systems) at the dilution of 1:10. The secondary antibody was the rabbit polyclonal anti-goat immunoglobulin antibody (bridge antibody; Jackson Immuno Research) diluted 1:250, and then with protein-A gold 5 nm (PAG5, CMC, Utrecht, The Netherlands) diluted in blocking solution 1:50 for 20 min at room temperature. Finally, the grids were stained for 10 min in 1.8% methyl cellulose plus 0.4% uranyl acetate, on ice. The grids were retrieved and after air-drying, they were examined under an electron microscope (FEI, Tecnai20; ThermoFisher Scientific, The Netherlands).

### Tissue processing and immunohistochemistry

All immunostainings of both WT and Gpr126^iECKO^ mice were carried out simultaneously and under the same conditions. Tissues were prepared and processed for immunohistochemical analysis as described previously (Bravi et al., 2015; Corada et al., 2013). Briefly, mice were anesthetized by intraperitoneal injection of Avertin 20 mg/kg (T48402, Sigma), and perfused with 1% PFA in PBS. The mouse brains were carefully dissected and post-fixed overnight by immersion in 4% PFA at 4 °C. The next day, they were washed in PBS and processed for sectioning as follows.

For vibratome sections, brains were embedded in 4% low-melting-point agarose, sectioned with a vibratome (100 mm thickness; VT1200s; Leica) and immune stained. Sagittal vibratome sections were incubated in PBST solution (PBS with 0.3% Triton X-100) supplemented with 5% donkey serum and containing the primary antibodies, overnight at 4 °C. This was followed by washes with PBST, followed by the secondary antibody solutions (overnight at 4 °C). Sections were then washed, post-fixed for 2 min in 1% PFA, and mounted in Vectashield that contained DAPI (H-1200; Vector). Detection of Claudin-5 (ab15106; Abcam) and Plvap (550563 clone MECA-32; BD Biosciences) was carried out as above, with the only exception that brains were post-fixed overnight in 100% methanol at 4 °C (instead of 4% PFA) and rehydrated for 1 h in PBS before sectioning.

For cryosectioning, fixed brains were cryo-protected in 30% sucrose overnight at 4 °C, embedded in Killik embedding medium (059801; BioOptica) and frozen on the surface of the cold isopentane/2-methylbutane next to the liquid nitrogen. Sections (10 μm) were cut using a cryostat (Leica), then mounted on positively charged glass slides and stained as described above.

To analyze the mouse retina vascular phenotype, the eyes were collected and fixed in 4% PFA in PBS for 2 h at 4 °C. Retinas were dissected as described previously (Corada et al., 2013) and incubated at 4 °C overnight in primary antibodies diluted in PBSTC buffer (PBS with 0.5% Triton X-100, 0.1 mM CaCl_2_) supplemented with 5% donkey serum, followed by the suitable species-specific Alexa Fluor-conjugated secondary antibody staining, overnight at 4 °C. These samples were then flat-mounted on glass microscope slides with Prolong Gold antifade reagent (P36930; ThermoFisher Scientific). For immunofluorescence analysis, negative controls using isotype matched normal IgG were included to check for antibody specificity.

### RNA interference

cBECs derived from WT pups at p18 were seeded into collagen I-coated wells (1.5 brain/10 cm^2^) and cultured in DMEM complete medium. After 3 days of puromycin selection, they were transfected with either non-targeting siRNAs (StealthTM RNAi Negative Control Duplexes; Invitrogen) or siRNAs against mouse Adgrg6 (MSS278013: #13 CGACUGCCAAGGGCCUGUCAUUUAA MSS210995: #95 GCCUCCAAAUUUGCUUGAGAAUUUA; MSS210997: #97 CCGUGUUACCCUAAUGACUACCCUA, ThermoFisher Scientific). Transfection was performed using Lipofectamine 2000 (LS11668019; ThermoFischer Scientific) or Lipofectamine RNAiMAX (13778150; ThermoFischer Scientific), according to the manufacturer instructions.

### Mouse ischemic stroke model

#### Proximal middle cerebral artery occlusion (MCAo)

For MCAo experiments, the *Cdh5(PAC)-CreER^T2^–*Gpr126^flox/flox^–Rosa26-Stopflox-EYFP mouse strain was used. Mice were injected from P1 to P4 and then the adult mice (8 weeks old) underwent 45-min left MCAo (Bacigaluppi et al., 2009). Briefly, the mice were anesthetized with 1.0% to 1.5% isofluorane (Merial, Assago, Italy) in 30% O_2_; their temperature was maintained between 36.0 °C and 36.5 °C, and Laser Doppler flowmetry (PeriFlux System 5000; Perimed) was used to monitor the procedure. Focal cerebral ischemia of the left MCA was induced with a 7-0 silicon-coated filament (#702034PK5Re; Doccol). The mice that successfully reached a stable cerebral blood flow below 30% of their initial baseline were included in the study, while the mice that experienced bleeding during surgery were excluded from the experiment. Laser Doppler recordings during surgery that had signal saturated, negative values or other technical registration problems (e.g., detachment of probe during ischemia) were excluded from the analysis of the laser Doppler flowmetry.

*Behavioral analysis:* The modified Neurological Severity Score (mNSS) grades neurological function was used, for a scale of 0 to 12 (normal score 0; maximal deficit score 12). mNSS is a composite of motor and balance tests (Bacigaluppi et al., 2009). Behavioral tests were performed by a researcher blinded to the treatment groups during the light phase of the circadian cycle, beginning 4 h after the lights were turned on.

#### Analysis of infarct size

On histological sections: For measurement of the ischemic lesion volume, 20-μm-thick coronal cryostat sections were sliced from 2 mm rostral to 4 mm caudal to the Bregma. One systematic random series of sections *per* mouse was stained with cresyl violet (Sigma), digitalized, and analysed with an Image J (NIH, USA) image analysis system. The lesion area was measured for each reference level by the ‘indirect method’, as described previously (Butti et al., 2012). For the representative three-dimensional volume rendering, representative brains images for each treatment group were stained by cresyl violet and traced onto images acquired by a color video camera mounted on a Leica microscope, using the Stereo Investigator v3.0 software (MicroBrightField, Inc., Colchester, VT) (Bacigaluppi et al., 2009; Butti et al., 2012; West et al., 1991).

On magnetic resonance imaging (MRI): MRI studies were performed on a 7 T preclinical magnetic resonance scanner (BioSpec 70/30 USR, Paravision 5.0; Bruker, Germany), equipped with 450/675 mT/m gradients (slew-rate: 3400–4500 T/m/s; rise-time: 140 μs). A phased-array rat-heart coil with four internal preamplifiers was used as receiver, coupled with a 72-mm linear-volume coil as transmitter. For the study, the mice were anesthetized with 1.5 to 2.0% isoflurane (Forane, Abbott), vaporized in 100% oxygen (flow: 1 L/min). For the measurement of lesion volume, RARE T2 sequences were acquired. Repetition time, 38 ms; echo time, 3200 ms; average number of, 12; total scan time, 10 min, 14 s; field of view: 14 mm × 15 mm; spatial resolution, 0.055 × 0.094 mm/pixel; matrix, 256 × 160 pixel. Volumes were calculated on coronal FLAIR images with the measurement tool of OsiriX Imaging Software: as volume of contralesional, right hemisphere without the volume of the right lateral ventricle (HVc); volume of the ipsilesional, left hemisphere subtracted from the volume of the left lateral ventricle (HVi); and uncorrected lesion volume (LVu) (Gerriets et al., 2004). The edema corrected lesion volume (HLVe) was then calculated as proportin (%) of the right hemisphere, using Equation (1):

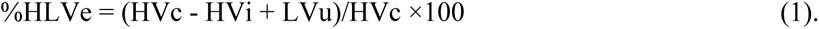

The proportion (%) of edema of the ischemic hemisphere HVc (HSE) was calculated as in Equation (2):

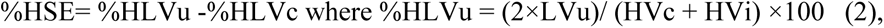

as described elsewhere (Gerriets et al., 2004).

*Tissue pathology*: At the indicated reperfusion times, the mice were deeply anesthetized and transcardially perfused with 25 mL PBS 0.1 M, pH 7.2 with EDTA, followed by 25 mL 4% PFA. For cryostat sectioning, the brains were carefully dissected, post-fixed overnight, cryoprotected in sucrose, and embedded in tissue-freezing medium. Frozen brains were cut into 20-μm-thick coronal cryostat sections.

The vessel density and the mean intensity of Pecam-1 in the contralateral and stroke region were quantified on the maximum projected image using a custom semi-automated Fiji plugin. The contralateral and stroke regions were drawn by hand and the vessel area was identified on the Pecam-1 channel by manual thresholding after rolling ball background subtraction and median filtering. For both regions, the vessel density was then evaluated as the ratio between the vessel area and the whole drawn region (stroke or contralateral), and the mean intensity of the signal in the vessel area was measured after rolling ball background subtraction.

For counting Iba-1- and CD68-positive cells in the contralateral and stroke regions, a custom semi-automated Fiji plugin identified and counted the cells within the drawn area using the ImageJ Find Maxima plugin, with the noise tolerance parameter set by hand.

#### Flow cytometry analysis

After 3 days, post-MCAo, mice were anesthetized, perfused with cold saline and the brain removed. The forebrain (separated from the bulbi, cerebellum, and brainstem) was divided into ipsilateral (ischemic) and contralateral hemispheres. Each hemisphere was processed separately. The brain tissues were cut with scissors into small pieces and incubated with 0.4 mg/mL collagenase type IV (C5138; Sigma-Aldrich) for 30 min at 37 °C. After the incubation, they were repeatedly passed through a syringe with a 19G needle, to obtain a homogeneous cell suspension, then filtered through a 70-μm mesh, suspended in PBS, and washed. Tissue homogenates were loaded onto an isotonic 90% Percoll (17089101; GE Heathcare) gradient for enrichment of CNS infiltrates (De Feo et al., 2017). Brain cell suspensions enriched on the Percoll gradient were then labeled and analyzed by flow cytometry (FACS Symphony; BD Biosciences) (Lelios and Greter, 2014), with data analysis by FlowJo software (Tree Star). The whole sample was processed. For brain analysis, cells were always gated on single and live cells by excluding dead cells, using the live/dead Zombie NIR Fixable Viability Kit (Cat. 423105; Biolegend). Neutrophils were identified as CD45^+^ CD11b^+^ Ly6G^+^ cells, and CD4^+^ and CD8^+^ T cells as CD45^+^ CD11b^−^ CD4^+^ and CD45^+^ CD11b^−^ CD8^+^ cells, respectively. B cells were identified as CD45^+^ CD11b^−^ B220^+^ cells, dendritic cells as CD45^+^ CD11b^+^ CD44^−^ CD11c^+^ MHCII^+^ cells, monocytes as CD45^+^ CD11b^+^ CD44^−^Ly6C^lo^ or Ly6C^hi^ CD11c^−^ MHCII^−^ cells, border-associated macrophages as CD45^+^ CD11b^lo^ CD44^+^ CD206^+^ cells, and microglia as CD45^+^ CD11b^+^ CD44^+^ cells.

### Western blotting

Total proteins were extracted by solubilizing the cells in boiling sample buffer (2×; 2.5% sodium dodecyl sulfate, 20% glycerol, 0.125 M Tris-HCl, pH 6.8) enriched in protease inhibitor R cocktail set III (EDTA free; 539134; Calbiochem) and phosphatase inhibitors (cocktail 2; P5726; and cocktail 3; P0044; Sigma-Aldrich). Lysates were incubated for 5 min at 100 °C, to promote protein denaturation, and then centrifuged at 16,000× *g* for 5 min, to pellet the cell debris. Protein concentrations were determined using BCA Protein Assay kits (23225; Pierce), according to the manufacturer instructions. Equal amounts of protein were loaded onto custom gels (Bio-Rad or Invitrogen), separated by SDS-PAGE, transferred to membranes (Protran nitrocellulose hybridisation transfer membrane; pore size, 0.2 μm; 10600001; GE Healthcare), and blocked for 1 h at room temperature in 5% milk or 5% bovine serum albumin, in 0.05% Tween-20 in Tris-buffered saline (TBS). The membranes were incubated overnight at 4 °C or for 1 h at room temperature, with the primary antibodies diluted in blocking solution. The membranes were rinsed three times with washing solution (0.05% Tween-20 in TBS) and incubated for 1 h at room temperature with horseradish-peroxidase-linked secondary antibodies (diluted in blocking solution). The specific binding was detected by Super Signal West Femto Maximum Sensitivity Substrate (34096; ThermoFisher Scientific) or Super Signal Dura Extended Duration Substrate (34076; ThermoFisher Scientific), using a ChemiDoc MP system (BioRad). The molecular masses of the proteins were estimated relative to the electrophoretic mobility of a co-transferred prestained protein marker (Precision Plus Protein Standards Dual Colour; 161-0374; BioRad). Densitometry analysis of the bands was carried out used the Fiji software (open source; htpp://fiji.sc/) and with the band analysis tools of the ImageLab software, version 4.1 (BioRad). GAPDH, tubulin, or VE-cadherin (loading control) were used to normalize the quantified bands, as specified in the Figures.

### Flow cytometry analysis

To obtain enriched fBECs by depletion of CD45-positive cells, the brains from 4 animals per group of WT and Gpr126^iECKO^ mice (P18) were processed as described above. Flow cytometry was performed using a Dako FACS instrument, Attune NxT flow cytometer (ThermoFisher Scientific). Phenotypic analysis was performed with the following antibodies: Rat anti-mouse active β1-integrin (active CD29 Clone 9EG7, 553715; BD Biosciences); Alexa647-Armenian hamster anti-mouse β1-integrin (total β1-integrin 102214; Biolegend); Alexa488-goat anti-mouse Pecam-1 (FAB3628G; R&D Systems). Fc receptor blocking reagent (130-092-575; Miltenyi) was used to block unwanted binding of antibodies to mouse cells expressing *Fc receptors.* Cy3 anti-rat secondary antibody (712-165-153; Jackson Immuno Research) was used to detect active β1-integrin. Cells were first gated to remove doublets, for light scatter and then for specific markers. Quantitative analysis was performed using Summit 4.2 from Dako, Kaluza software from Beckman Coulter. Alexa488 conjugated anti-PECAM1 antibody was used to identify the EC population, where the proportions (%) of positive cells expressing active and total β1-integrin were analyzed.

### Sprouting angiogenesis assay

For the angiogenic sprouting assay (Heiss et al., 2015), WT and Gpr126^iECKO^ ECs were isolated from brains of pups at P18, as previously described. Briefly, after dissociation, myelin cell debris was removed according to the manufacturer protocol. ECs were enriched by depletion of CD45-positive cells with CD45 microbeads (30-052-301; Miltenyi Biotech), followed by positive selection using CD31 microbeads (30-097-418; Miltenyi Biotech). Then, the enriched ECs were resuspended in MCDB-131 complete medium containing 0.1% methylcellulose (4000 cP, #M0512; Sigma). For spheroid formation, 10000 cells per 25 µL methylcellulose medium were seeded in wells of U-bottomed 96-wells suspension plates and incubated for 5 days. Then, spheroids (25 µL) were collected and resuspended in 600 µL/well 3 mg/mL Type I rat tail collagen (High Conc collagen I, 354249; Corning) in 199 medium (21180021; ThermoFisher Scientific) and placed at 37 °C for 30 min. After polymerization of the collagen gel, the spheroids were stimulated with 80 ng/mL mVEGF (450-32; Peprotech) and 50 ng/mL b-FGF (450-33; Peprotech), to induce sprouting, for 48 h in complete medium. Pictures were acquired using the Thunder imaging system and the 20× objective. Sprouting numbers and lengths were analyzed using the line tool of Fiji.

### Scratch wound migration assay

cBECs were cultured on 6-well plates coated with collagen I. After reaching confluency, the monolayers were scratched cross-wise with a p200 pipette tip, washed with MCDB-131 medium and 2% fetal bovine serum, and imaged at time 0 using an Optika Microscopy C-P6 FL imaging system and the 20× objective. Then the 6-well plates were incubated for 36 h and were imaged at the end point. The wound width at T0 (scratch time) and after 36 h was estimated as the average distance of three vertical lines drawn by hand between the two cell layers and always at the same positions along the wound, using the line tool of Fiji.The wound closure was then calculated, as normalizing using the T0 measurements.

### Co-immunoprecipitation assay

Confluent monolayers of iBECs transfected with GFP and Gpr126-myc (2 ×10^7^ cells) were solubilised in cold lysis CHAPS buffer (50 mM Tris-HCl, pH 7.4, 50 mM NaCl, 2 mM MgCl_2_, 1 mM EDTA, 15 mM CHAPS (C5070; Sigma), cOmplete Protease Inhibitor Cocktail (4693116001; Roche) and phosphatase inhibitors (P2850, P5726; Sigma-Aldrich), and incubated on ice for 1 h. For Gpr126 immunoprecipitation, after preclearing with protein A-Sepharose, 4.5 mg of the protein extracts were incubated with anti-c-myc agarose affinity gels, antibody rabbit (A7470, Sigma) overnight at 4 °C. For β1-integrin immunoprecipitation, after preclearing with protein G-Sepharose 4B plus, 9 mg protein extracts were incubated with goat anti-β1-integrin (AF2405; R&D Systems) overnight at 4 °C, followed by incubation with protein G-Sepharose 4B plus for 1 h. The immune complexes were eluted in 100 μL elution buffer (8 M urea in 100 mM Tris-HCl, pH 7) and Western blotting samples were prepared by adding sample buffer (3× Laemmli Sample Buffer 1610747; BioRad) plus 3.57 M β-mercaptoethanol (2-mercaptoethanol, 63689; Sigma-Aldrich). The following antibodies were used: anti-Gpr126 rabbit (Ab75456; Abcam), anti-myc mouse (2276; Cell Signaling), anti-Lrp1 mouse (ab215997; Abcam), anti-β1-integrin rabbit (4706; Cell Signaling), anti-α1 integrin goat (ab243032; Abcam), and anti-integrin α3 goat (AF2787; R&D).

### Statistical analysis

The normality of the datasets was assessed using Shapiro-Wilk normality tests. For datasets with normal distributions, two-sided unpaired Welch’s t-tests (for pairwise comparison) or one-way Brown-Forsythe ANOVA followed by Dunnett’s T3 tests (for *post-hoc* multiple comparisons) or two-way ANOVA were used. Non-parametric (Mann-Whitney) tests were applied to datasets that did not show normal distributions. Wherever applicable, the details about the statistical test applied are specified in the Figure legend. The standard software package GraphPad Prism (v 9.4.0) was used.

### Image acquisition, processing, and analysis

Confocal microscopy was performed using a confocal microscope (SP8; Leica). For image analysis, the Fiji software was used (open source, http://fiji.sc/). The figures were assembled and processed using Adobe Photoshop and Adobe Illustrator. The only adjustments used in the preparation of the figures were for brightness and contrast. For comparison purposes, different sample images of the same antigen were acquired under constant acquisition settings.

To quantify Gpr126 mRNA expression levels using RNA scope in the brain microvasculature, an automated custom Fiji plugin (Schindelin et al., 2012) was developed. The number of Gpr126 RNA scope spots in the vessels assessed by Claudin-5 positive regions was measured on the maximum projected images. Claudin-5 regions were identified on the 488-channel using the Li’s threshold (ttps://imagej.net/plugins/auto-threshold#li) method after maximum filtering, rolling ball background subtraction and Gaussian filtering. Within these regions the number of Gpr126 RNA scope spots were identified using the Find Maxima tool after rolling ball subtraction on the 647-channel.

Cadaverine leakage was measured as follows. Each mouse brain sagittal section was acquired under 10× magnification. Then, using Fiji, eight regions of interest (ROIs) were defined (three in the cortex, one in the olfactory bulb, cerebellum, midbrain, pons, medulla, hypothalamus, striatum). The ROI sizes were defined to roughly cover 80% of the area of the region. The mean intensities of each ROI were measured. The size and the relative position of the ROIs were kept constant across each comparison. Two sections for each sample were used for quantification. Quantification of the leakage was carried out using two distinct operators. The first operator acquired the confocal stacks, while the second operator performed the quantification in a blind test.

Quantification of brain vessel average width was determined using two custom Fiji plugins. The first plugin generated a distance map as an intermediate result, using the local thickness tool after the Huang thresholding method on the image filtered with a median and Gaussian filter, in which the vessels where stained for podocalyxin. The second plugin applied a grid on the local thickness distance map and for each tile determined the vessel average width as the mean intensity of the pixels with a value above 0 (where intensities represent the distances on the local thickness distance map).

The numbers of microcapillaries in the brains stained with Pecam-1 were assessed using a custom Fiji plugin. This tool identified and counted the microcapillaries using the triangle thresholding method on the sum projection of the Pecam-1 channel filtered with a Median filter.

The vessel (podocalyxin, Pecam-1) coverage by pericyte (Pdgfrβ, CD13) and BM (fibronectin, collagen IV, laminin α2) was evaluated using a semi-automated Fiji plugin on the maximum projected image. Vessel region was identified by manual thresholding after rolling ball background subtraction, maximum and median filtering. Within this region, the coverage area (pericytes or BM) was quantified by manual thresholding after rolling ball background subtraction and Gaussian filtering.

Plvap and claudin-5 mean intensities in the vessel area (assessed by podocalyxin) were quantified using a custom Fiji plugin. This tool identified the vessel areas using the triangle thresholding method on podocalyxin channel filtered with a median and maximum filter after rolling ball background subtraction. Within the vessel area, the claudin-5 intensity is measured after background correction, which was estimated using the mean intensity of a region drawn by hand.

Quantification of tip cells in the retina was carried out by two separate operators. The first operator acquired the confocal stacks, and the second operator manually counted tip cells in a blind test. Cell counting was through the cell counter plugin of Fiji.

The radial expansion in the retina was measured as the ratio between the mean distance covered by the vessels (growing from the optic nerve) and the total length of the petal. The lengths were measured by hand using the line tool of Fiji on the maximum projected image.

The vessel density in the retina was determined using a semi-automated Fiji plugin. This tool quantified the vessel density as the ratio between the area covered by vessels (colored pixels of stained vessels) and the whole retina region (dark pixels of the retina tissue) on the maximum projected image. The whole retina region area was identified by manual thresholding and Fill Holes binary operation after a large Gaussian filtering. Within this region, the vessel area was identified by manual thresholding after median filtering.

Quantification of the Edu EC nuclei in the retina was carried out on the maximum projected image using a custom automated Fiji plugin. The vessel area (CD93 staining) was defined using the Li thresholding method after filtering the image with a median filter. Within this area the plugin identified the nuclei EC area (Erg staining) using the Otsu thresholding method, after filtering the image with median and Gaussian filters. The double Erg- and Edu-positive nuclei were counted with the ImageJ Find Maxima tool within the vessel area.

To measure the ratio between the Ki67- or Brdu-positive cells and Erg-positive cBECs, a custom Fiji plugin was developed. The plugin counted the Erg-positive nuclei using the Yen’s threshold method on the sum projected image, after median and Gaussian filtering. Within the Erg-positive nuclei, the plugin counted the cells on the Ki67 and Brdu staining using the same process.

### Antibodies

The antibodies used in this study were: anti-Gpr126 rabbit (1:500, ab75456; Abcam; WB; IF), anti-Gpr126 rabbit (1:10, Ab218046; Abcam; EM) anti-Gapdh mouse (1:1000, sc32233; Santa Cruz; WB), anti-podocalyxin goat (1:200, AF1556; R&D; IF), anti-tubulin mouse (1:2000, T9026; Sigma; WB), anti-phospho-Creb rabbit (1:1000, 9198; Cell Signaling; WB), anti-Creb rabbit (1:1000, 4820; Cell Signaling; WB), anti-myc mouse (1:1000, 2276; Cell Signaling; WB), anti-c-myc agarose affinity gel antibody rabbit (A7470; Sigma; IP), anti-Lrp1 mouse (1:1000; ab215997; Abcam; WB), anti-integrin β1 goat (1:200; AF2405; R&D; IP), anti-integrin β1 rabbit (1:1000; 4706; Cell Signaling; WB), rat anti-mouse β1-integrin (1:50, 553715; BD Biosciences, active β1, FACS); alexa Fluor 647 anti-mouse/rat β1-integrin (1:50, 102214; Biolegend, FACS); anti-integrin β1 (1:10, AF 2325; R&D Systems; EM), anti-integrin α1 goat (1:500; ab243032; Abcam; WB), anti-integrin α3 goat (1:3500; AF2787; R&D; WB), Horse anti-rabbit biotinylated IgG (1:400; VC-BA-1100-MM15; Vector Laboratories; IF), streptavidin Alexa Fluor 555 conjugate (1:400; S32355; Invitrogen; IF), anti-GFP chicken (1:1000; ab13970; Abcam; IF), anti-Pecam-1 goat (1:200; AF3628; R&D Systems; IF), and anti-isolectin-B4 (1:200; B-1205; Vector Laboratories; IF); anti-m/rCD31/PECAM-1 Alexa Fluor 488 Conjugated (1:50, FAB3628G; R&D Systems); Claudin-5 (1:200, Ab15106; Abcam; IHC) and Plvap (1:200, 550563 clone MECA-32; BD Biosciences, IHC); BUV805 Rat anti-Mouse CD3 molecular complex (Monoclonal; clone 17A2; BD Biosciences;1:200; Cat#741982, FACS). APC Rat anti mouse CD4 (Monoclonal; clone RM4-5; BD Biosciences; 1:300; Cat#553041, FACS). PE-Cy7 Rat anti-mouse CD8 (Monoclonal; clone 53-6.7, FACS); BD Biosciences; 1:300; Cat#561097. BUV737 Rat anti-Mouse CD11b (Monoclonal; clone M1/70; BD Biosciences; 1:200; Cat#612800, FACS). PE-Cy5.5 Armenian Hamster anti-Mouse CD11c (Monoclonal; clone N418; BD Bioscience; 1:400; Cat#35-0114-82, FACS). PE Rat anti-Mouse/Human CD44 (Monoclonal; clone IM7; Biolegend; 1:100; Cat#103007, FACS). BUV395 Rat anti-Mouse CD45 (Monoclonal; clone 30-F11; BD Biosciences; 1:400; Cat#564279, FACS). BUV661 Rat anti-Mouse CD45R/B220 (Monoclonal; clone RA3-6B2; BD Biosciences; 1:400; Cat#612972, FACS). BV711 Mouse anti mouse CD64 (Monoclonal; clone X54-5/7.1; Biolegend; 1:100; Cat#139311, FACS). BV421 Rat anti mouse CD68 (Monoclonal; clone FA-11; Biolegend; 1:300; Cat#137017, FACS). AlexaFluor700 Rat anti mouse CD206 (Monoclonal; clone C068C2; Biolegend; 1:100; Cat#141734, FACS). PE/Dazzle™ 594 Mouse anti-mouse CX3CR1 (Monoclonal; clone SA011F11; Biolegend; 1:300; Cat#149013, FACS). Biotin Rat anti-Mouse F4/80 (Monoclonal; clone Cl:A3-1; Bio Rad; 1:100; Cat# MCA497BB, FACS). BV605 Rat anti-Mouse Ly6C (Monoclonal; clone HK1.4; Biolegend; 1:400; Cat#128036, FACS). BUV563 Rat anti-Mouse Ly6G (Monoclonal; clone 1A8; BD Biosciences; 1:200; Cat#612921, FACS). BB700 Rat anti-Mouse MHCII (Monoclonal; clone M5/114.15.2; BD Biosciences;1:400; Cat#746197, FACS). BV786 Mouse anti-Mouse NK1.1 (Monoclonal; clone PK136; BD Biosciences; 1:100; Cat#740853, FACS). Zombie NIR™ Fixable Viability Kit; Biolegend; 1:500; Cat#423106, FACS). Streptavidin BV650 (Biolegend; 1:400; Cat#405231, FACS). HRP-linked anti-mouse and anti-rabbit (1:2000; Cell Signaling; WB), AlexaFluor 555-conjugated donkey anti-rabbit (1:400; Invitrogen; IF); AlexaFluor 488-conjugated donkey anti-goat (1:400; Invitrogen; IF); Cy3-donkey anti-rat IgG (1:200, Jackson Secondary Antibody, 712-165-153; FACS).

## Non-standard abbreviations and acronyms

AU: arbitrary units
BBB: blood-brain barrier
BM: basement membrane
cAMP: cyclic adenosine monophosphate (or 3’,5’-cyclic adenosine monophosphate)
cBECs: cultured murine endothelial cells derived from the brain microvasculature
cLEC: cultured lung-derived endothelial cells
CNS: central nervous system
CREB: cAMP response-element binding protein
ECM: extracellular matrix
ECs: endothelial cells
eGFP: enhanced green fluorescent protein
fBECs: freshly isolated brain endothelial cells
fLECs: freshly isolated lung endothelial cells
GAIN: GPCR autoproteolysis-inducing domain
GO: gene ontology
GPCRs: G-protein–coupled receptors
JAM-A: junctional adhesion molecule-A
MCAo: middle cerebral artery occlusion
NS: not statistically significant
NVU: neurovascular unit
PECAM-1: platelet/endothelial cell adhesion molecule-1
RT-qPCR: real time quantitative polymerase chain reaction
TJs: tight junctions
VEC: vascular endothelial cadherin
WT: wild-type

## Acknowledgements

We thank Stefan Lieber and Lena Claesson-Welsh for their valuable insights. We thank Christopher P. Berrie for editorial assistance and Giampietro Costanza for critical reading of the manuscript. We are indebted to Steffen E Storck for providing the primary brain endothelial cells from Slco1c1-Cre-ER^T2^/Lrp1^flox/flox^ mice.

M.G., N.K conceived and designed the study; N.K., M.G., A.A.S. performed the *in-vitro* experiments; C.M. and N.K. performed the *in-vivo* experiments and analyzed the results with M.G.; E.M. and S.M performed imaging analysis. M.G.L., N.K., E.D. G.M. and C.C. contributed to scientific discussions; F.I. and F.Z. performed the bioinformatic analyses; K.B. and A.P. performed the junctional image analysis; M.B. and G.S.G. conducted the stroke experiments and analyzed the data with M.G.; N.K., GV.B. and A.A.M. performed the electron microscopy experiments; M.G. wrote the manuscript, which was revised by C.C, E.D. and M.B. The final form of the manuscript has been approved by all of the authors.

## Supplementary figures legends

**Supplementary Figure 1.**
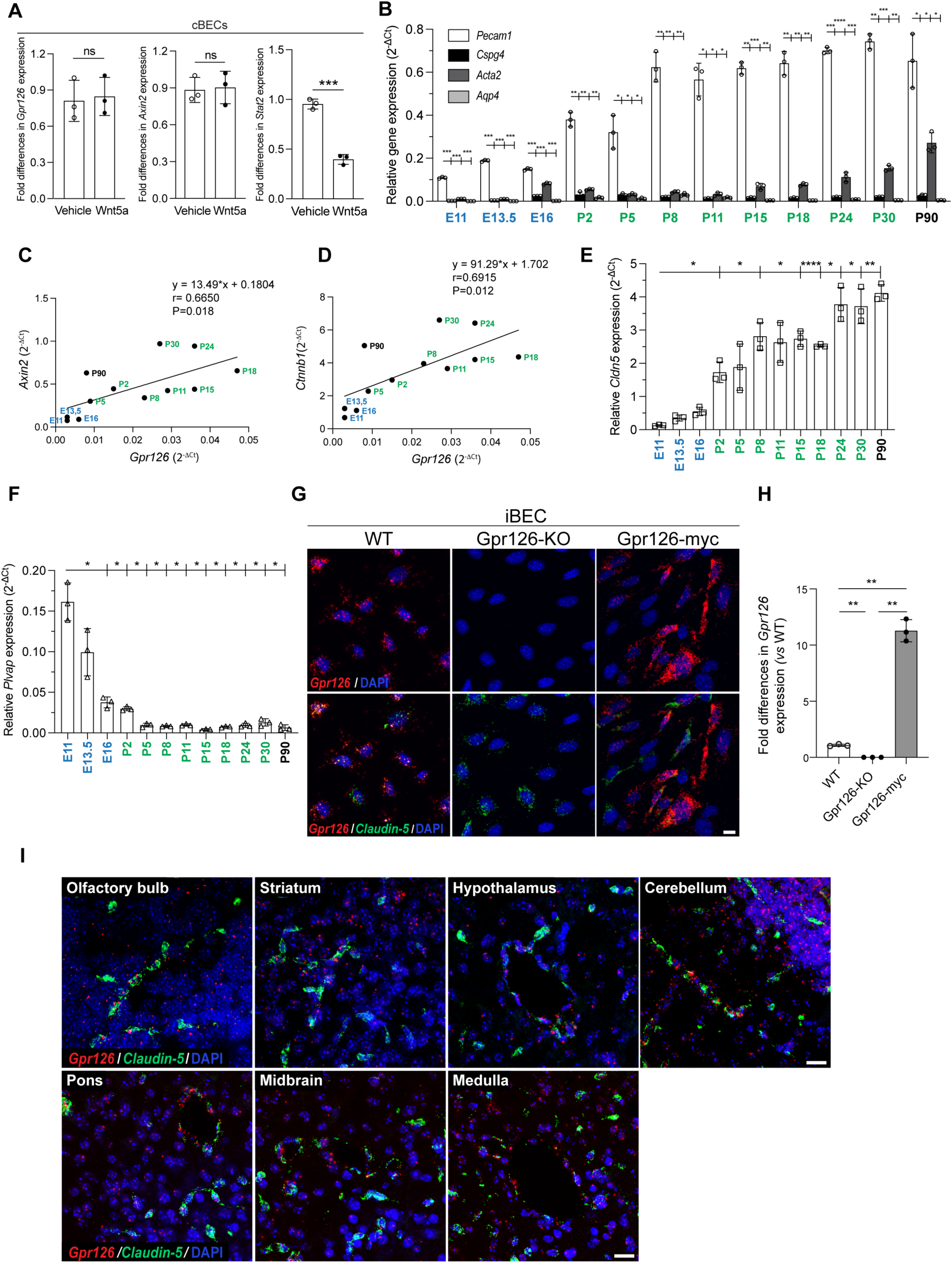
Gpr126 is a target of the canonical Wnt signaling and is expressed in the brain microvasculature during BBB development. **A,** Quantification of fold differences in *Gpr126* and *Axin2* and *Stat2* gene expression in cBECs from adult WT mice following RT-qPCR analysis. The cells were treated with recombinant Wnt5a or vehicle as control. Each symbol represents a single experiment (n=4 WT mice for each independent experiment, as means ±SD). ***, *P*<0.0005; ns=not significant (unpaired t-tests with Welch’s correction). **B,** Quantification of relative expression of *Pecam1* (ECs), *Cspg4* (pericytes), *Acta2* (smooth muscle cells) and *Aqp4* (astrocytes) in freshly isolated brain endothelial cells (fBECs) from WT mice at different stages during embryonic (E11-E16), postnatal (P2-P30) development, and in the adult (P90) following RT-qPCR analysis. Each symbol represents a single experiment (n=8 mice for embryos, n=5 mice for P2-P5 and n=3 mice for P8-P90 for each independent experiment, as means ±SD). \**P*<0.05; \*\**P*<0.005; \*\*\**P*<0.0005 (Brown-Forsythe and Welch ANOVA, Dunnett’s T3 multiple comparisons tests). **C, D,** Pearsońs correlation analysis (r) of gene expression in fBECs from WT mice at different stages during embryonic (E11-E16), postnatal (P2-P30) development and in the adult (P90) following RT-qPCR analysis. **C,** Positive correlation between *Adgrg6* expression and *Axin2* (p = 0.018, r = 0.6650). **D,** Positive correlation between *Adgrg6* expression and *Ctnnb1 (*p = 0.012, r = 0.6915). **E,** Quantification of relative *Cldn5* expression in fBECs from WT mice at different stages during embryonic (E11-E16), postnatal (P2-P30) development and in the adult (P90) following RT-qPCR analysis. Each symbol represents a single experiment (n=8 mice for embryos, n=5 mice for P2-P5 and n=3 mice for all the rest time points for each independent experiment, as means ±SD). \**P*<0.05; \*\**P*<0.005; ****, *P*<0.00005 (Brown-Forsythe and Welch ANOVA, Dunnett’s T3 multiple comparisons tests). **F,** Quantification of relative *Plvap* expression in fBECs from WT mice at different stages during embryonic (E11-E16), postnatal (P2-P30) development and in the adult (P90) following RT-qPCR analysis. Each symbol represents a single experiment (n=8 mice for embryos, n=5 mice for P2-P5 and n=3 mice for all the rest time points for each independent experiment, as means ±SD). \**P*<0.05 (Brown-Forsythe and Welch ANOVA, Dunnett’s T3 multiple comparisons tests). **G,** Representative confocal images of RNA scope *in-situ* hybridization for *Gpr126* (red) and *Claudin-5* (green) mRNA in immortalized brain endothelial cells (iBECs) expressing endogenous Gpr126, without Gpr126 (Gpr126-KO) and overexpressing myc-tagged Gpr126 (Gpr126-myc). DAPI (4′,6-diamidino-2-phenylindole; blue) stains nuclei. Scale bar: 10 μm. **H,** Quantification of fold differences in *Gpr126* expression in iBECs, as shown in G, following RT-qPCR analysis. Each symbol represents a single experiment (n=3, as means ±SD). \*\**P*<0.005 (Brown-Forsythe and Welch ANOVA, Dunnett’s T3 multiple comparisons tests). **I,** Representative confocal images of RNA scope *in-situ* hybridization for *Gpr126* (red) and *Claudin-5* (green) mRNA in different regions of mouse brain at P18. DAPI (blue) stains nuclei. Scale bar: 20 μm.

**Supplementary Figure 2.**
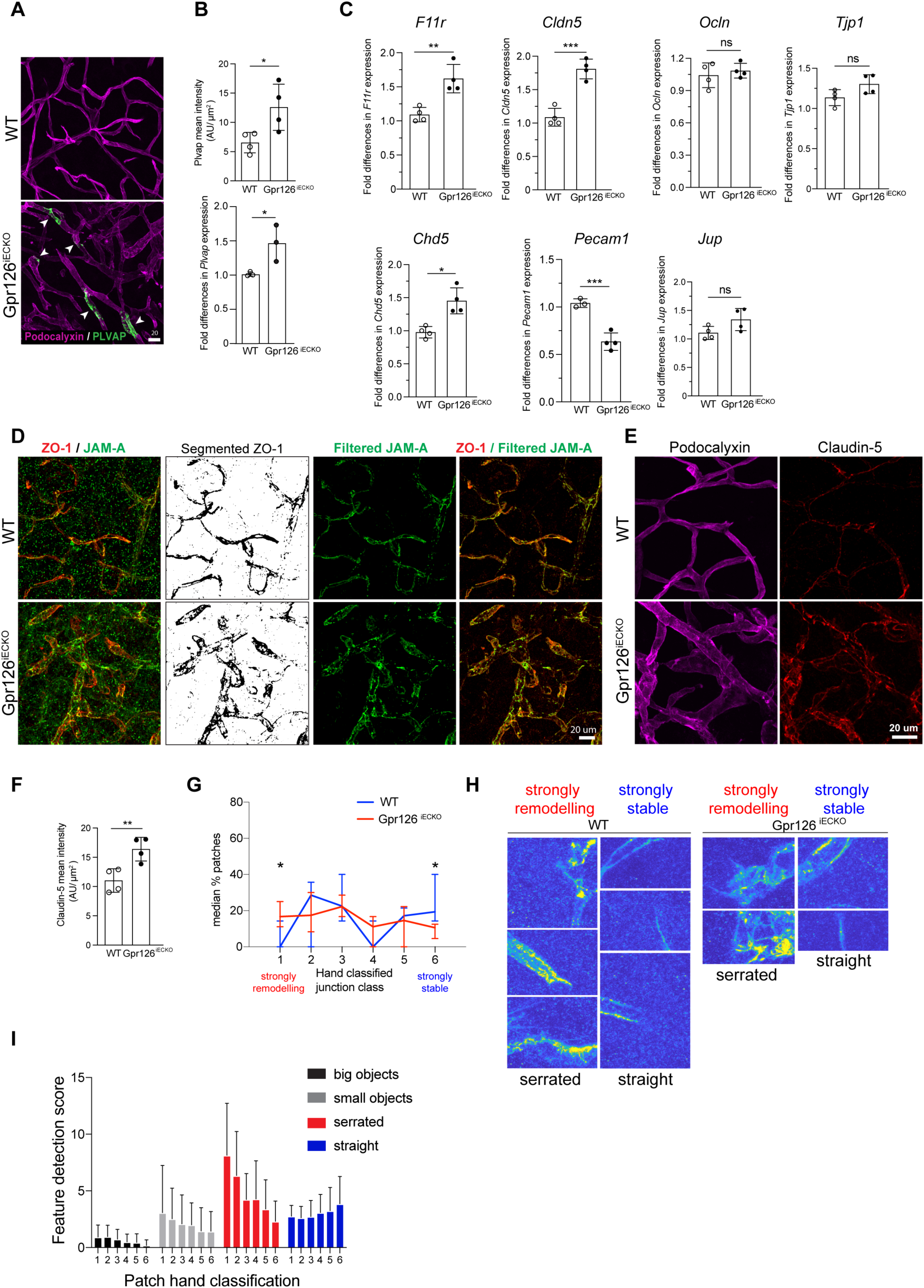
Endothelial junction organization is negatively affected in Gpr126iECKO mice and contributes to the loss of BBB function. **A,** Representative confocal images for podocalyxin (ECs, magenta) and plvap (fenestrated vessels, green) in brain cortex vibratome sections (100 μm) of WT and Gpr126^iECKO^ mice at P18. The arrowheads indicate the presence of Plvap staining. Scale bar: 20 μm. **B,** Quantification of Plvap signal (mean intensity) in the vessels area of brain cortex cryosections, as shown in **B** and expressed as arbitrary units (AU); Each symbol represents a mean of 6 sections for each mouse (n=4 WT and 4 Gpr126^iECKO^ mice, as means ±SD). *, *P* ≤0.05 (unpaired t-tests with Welch’s correction). Bottom: quantification of fold differences in *Plvap* gene expression in fBECs from WT and Gpr126^iECKO^ pups at P18. Each symbol represents a mouse. (n=3 WT and 3 Gpr126^iECKO^ mice, as means ±SD). *, *P* ≤0.05 (unpaired t-tests with Welch’s correction). **C,** Quantification of the fold differences in *F11 receptor* (*F11r*), *Claudin-5* (*Cldn5*), *Occludin* (*Ocln*), *Tight junction protein 1* (*Tjp1*), *Ve-cadherin* (*Cdh5*), *Platelet and endothelial cell adhesion molecule-1* (*Pecam1*) and *Junction plakoglobin* (*Jup*) gene expression in fBECs from WT and Gpr126^iECKO^ pups at P18. Each symbol represents a mouse (n=3-5 WT and 3-5 Gpr126^iECKO^ mice, as means ±SD). *, *P*<0.05; **, *P*<0.005; ***, *P*<0.005; ns=not significant (unpaired t-tests with Welch’s correction). **D,** Representative confocal images for zonula occludens-1 (ZO-1; red) and Junctional adhesion molecule-A (JAM-A; green) in brain cortex vibratome sections (100 μm) of WT and Gpr126^iECKO^ mice at P18. ZO-1 staining was segmented with threshold 350-4.096 (Segmented ZO-1) for merged images of filtered JAM-A (ZO-1/filtered JAM-A) Scale bar: 20μm. (n=3, WT and n=3, Gpr126^iECKO^). **E,** Representative confocal images for podocalyxin (magenta) and claudin-5 (red) in brain cortex vibratome sections (100 μm) of WT and Gpr126^iECKO^ mice at P18. Scale bar: 20 μm. **F,** Quantification of claudin-5 signal (mean intensity) in the vessels area of brain cortex cryosections, as shown in E and expressed as arbitrary units (AU); Each symbol represents a mean of 6 sections for each mouse (n=4 WT and 4 Gpr126^iECKO^ mice, as means ±SD). *, *P* ≤0.05 (unpaired t-tests with Welch’s correction). **G,** Quantification of claudin-5 patterning between strongly remodeling (1) to strongly stable (6) junctional shapes using the open-source Patching software (Bentley et al., 2014) for WT and Gpr126^iECKO^ mice (slices where there are > 4 non-empty patches included only, as slices with fewer than this were too sparse to analyse). (n=14 WT, n=18 Gpr126^iECKO^ images, as means ±SE). *, *P* ≤0.005 (one-sided Wilcoxon rank-sum test for equal medians). **H,** Randomly chosen patches from WT and Gpr126^iECKO^ images hand classified as strongly remodeling or stable. **G,** Automatic feature detection scores per hand classified patch class over all images analyzed. Scores were calculated using the automatic image analysis segmentation and object quantification functions for “serratedness” and straightness as defined in (Bentley et al., 2014)

**Supplementary Figure 3.**
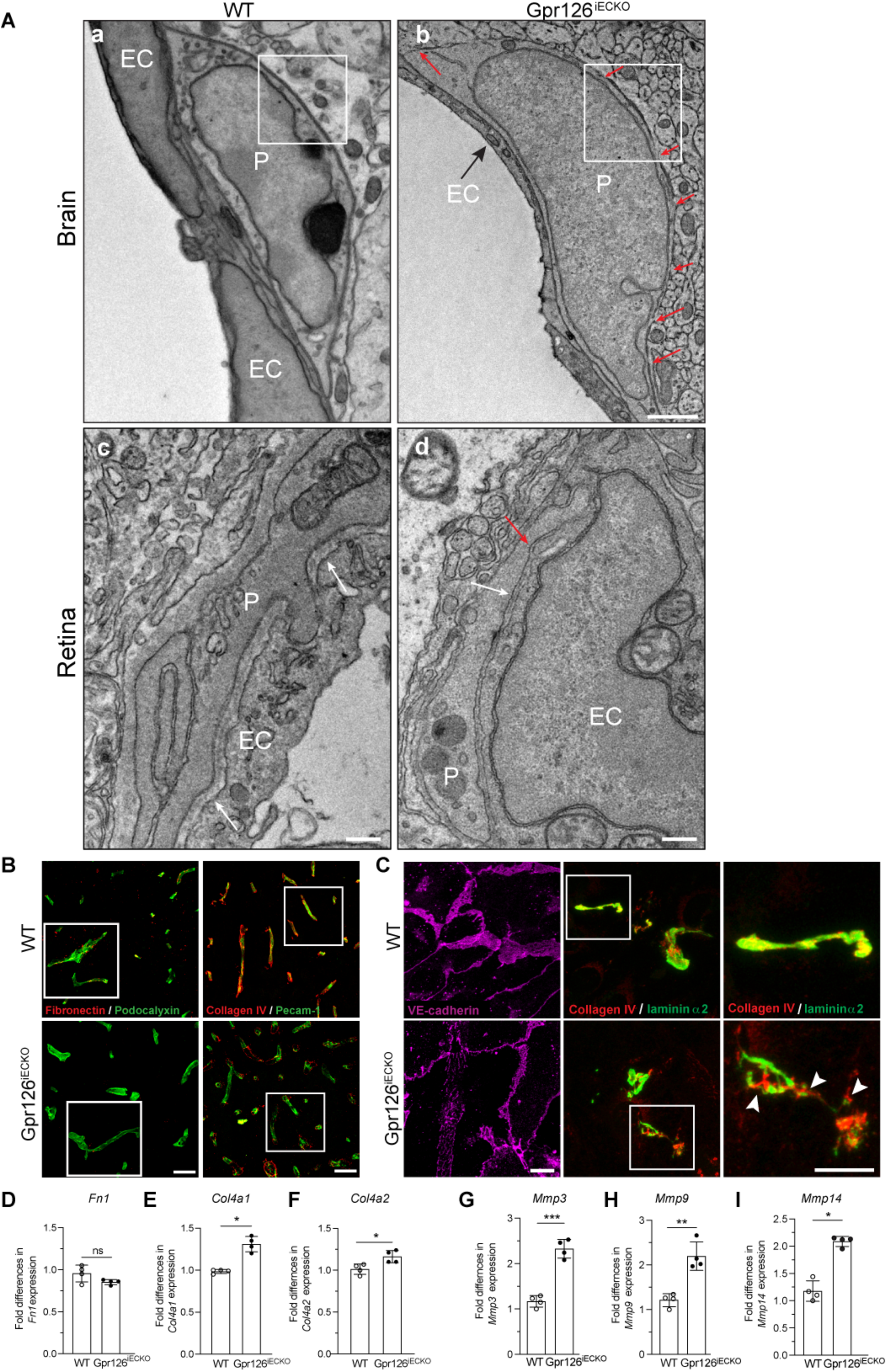
Gpr126 is required for proper BM protein deposition. **A,** Representative tomography virtual slices of brain (a and b) and retina (c and d) blood capillary longitudinal cortex section from WT and Gpr126^iECKO^ mice at P18. White arrows indicate regular thickness of the BM. Red arrows indicate decreased thickness or absence of the BM. White boxes show regions that are magnified in Figure 3 A. EC= endothelial cell; P=pericyte. Scale bar (a, b): 1 μm. Scale bar (b, c): 330 nm. **B,** Representative confocal images of brain cortex cryosections (4 μm) from WT and Gpr126^iECKO^ mice at P18. For the vasculature, podocalyxin (green), or Pecam-1 (green), were used. The BM was stained for Fibronectin (red) or Collagen IV (red). Merged images are shown. White boxes show magnified regions of Figure 3 A. Scale bar: 200 μm. **C,** Representative confocal images for VE-cadherin (magenta), collagen IV (red) and laminin α2 (green) in cultured brain-derived endothelial cells (cBECs) from WT and Gpr126^iECKO^ mice at P18. White boxes show magnified region depicted on the right. Collagen IV/Laminin α2: arrowheads show the absence of colocalization of the two proteins. Scale bar: 10 μm. **D-I**, Quantification of fold differences in *Fn1* (*Fibronectin*), *Collagen4a1* (*Col4a1*), *Collagen4a2* (*Col4a2*), *Matrix metalloprotease 3* (*Mmp3*), *Matrix metalloprotease 9* (*Mmp9*) and *Matrix metalloprotease 14* (*Mmp14*) gene expression in fBECs from WT and Gpr126^iECKO^ mice at P18. Each symbol represents a mouse (n=4 WT and 4 Gpr126^iECKO^ mice, as means ±SD). *, *P*<0.05; **, *P*<0.005; ***, *P*<0.005; ns=not significant (unpaired t-tests with Welch’s correction).

**Supplementary Figure 4.**
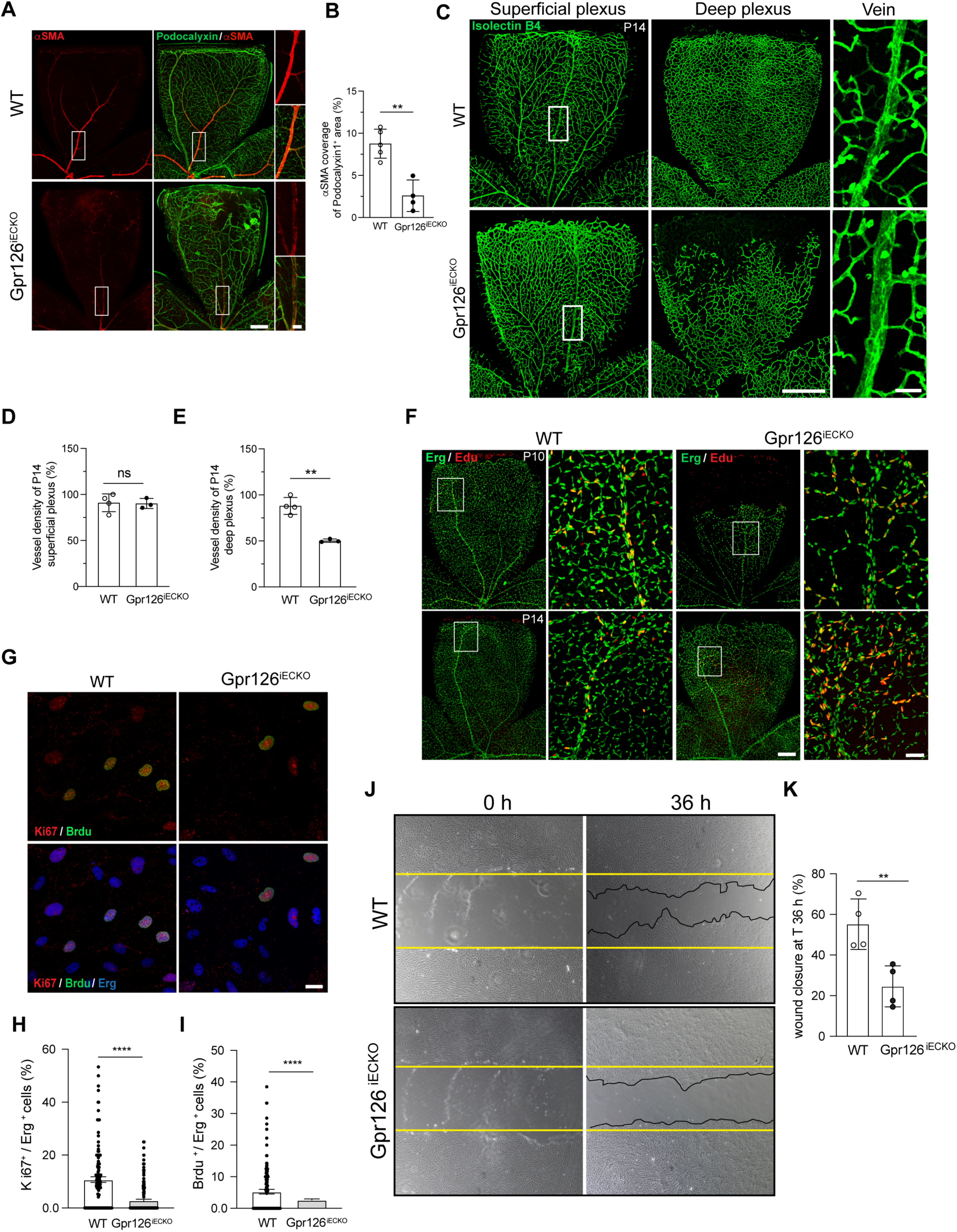
Gpr126 is required for the correct development of the retinal vasculature that supports EC proliferation and migration. **A,** Representative confocal images for Asma (A-smooth muscle actin, veins; red) and podocalyxin (green) in WT and Gpr126^iECKO^ mouse retinas at P18. Scale bar: 500 μm. White rectangles are shown in magnification on the right of each image. Scale bar: 100 μm. **B,** Quantification of Asma coverage of podocalyxin-positive area, expressed as percentage, as shown in A. Each symbol represents a retina for each mouse. (n=5 WT and 4 Gpr126^iECKO^ mice, as means ±SD). **, *P*<0.005 (unpaired t-tests with Welch’s correction). **C,** Representative confocal images for isolectin B4 (green) in the superficial or the deep retinal vasculature plexus from WT and Gpr126^iECKO^ mice at P14. Scale bar: 500 μm. White boxes are shown in magnification on the right of each image. Scale bar: 50 μm. **D-E,** Quantification of the vessel density superficial (**D**) and deep (**E**) plexus of the retinas, as shown in **C,** expressed as percentage. Each symbol represents a retina for each mouse. (n=4 WT and 3 Gpr126^iECKO^ mice, as means ±SD). **, *P*<0.005; ns=not significant (unpaired t-tests with Welch’s correction). **F,** Representative confocal images for Erg (EC nuclei, green) and Edu (proliferating cell nuclei, red) in WT and Gpr126^iECKO^ mouse retinas at P10 (upper panels) and P14 (lower panels). Scale bar: 500 μm. White rectangles are shown in magnification on the right of each image. Scale bar: 100 μm. **G,** Representative confocal images for Ki67 (proliferating cells, red), Brdu (S phase cells, green) and Erg (EC nuclei, blue) in cBECs from WT and Gpr126^iECKO^ mice at P18. Scale bar: 20 μm. **H-I,** Quantification of the percentage ratio of cells positive for Ki67 (**H**) or Brdu (**I**) to the total number of cells (Erg-positive), as shown in **G**. Each symbol represents a mean of 24 fields for each mouse (n=4 WT and 4 Gpr126^iECKO^ mice, as means ±SEM). \*\*\*\**P*<0.00005 (unpaired t-tests with Welch’s correction). **J,** Representative phase-contrast images of the scratch wound migration assay at time 0 and 36 h post-scratch performed in cBECs from WT and Gpr126^iECKO^ mice at P18. The yellow lines indicate the edges of the wound at time 0. The punctuated black lines indicate the boundaries of the wound after 36 h. **K,** Quantification of the wound closure after 36h to the total width of the wound at time 0, expressed as a percentage. Each symbol represents a single experiment (n=20 WT and 20 Gpr126^iECKO^ mice for each independent experiment, as means ±SD). \*\**P*<0.005 (unpaired t-tests with Welch’s correction).

**Supplementary Figure 5.**
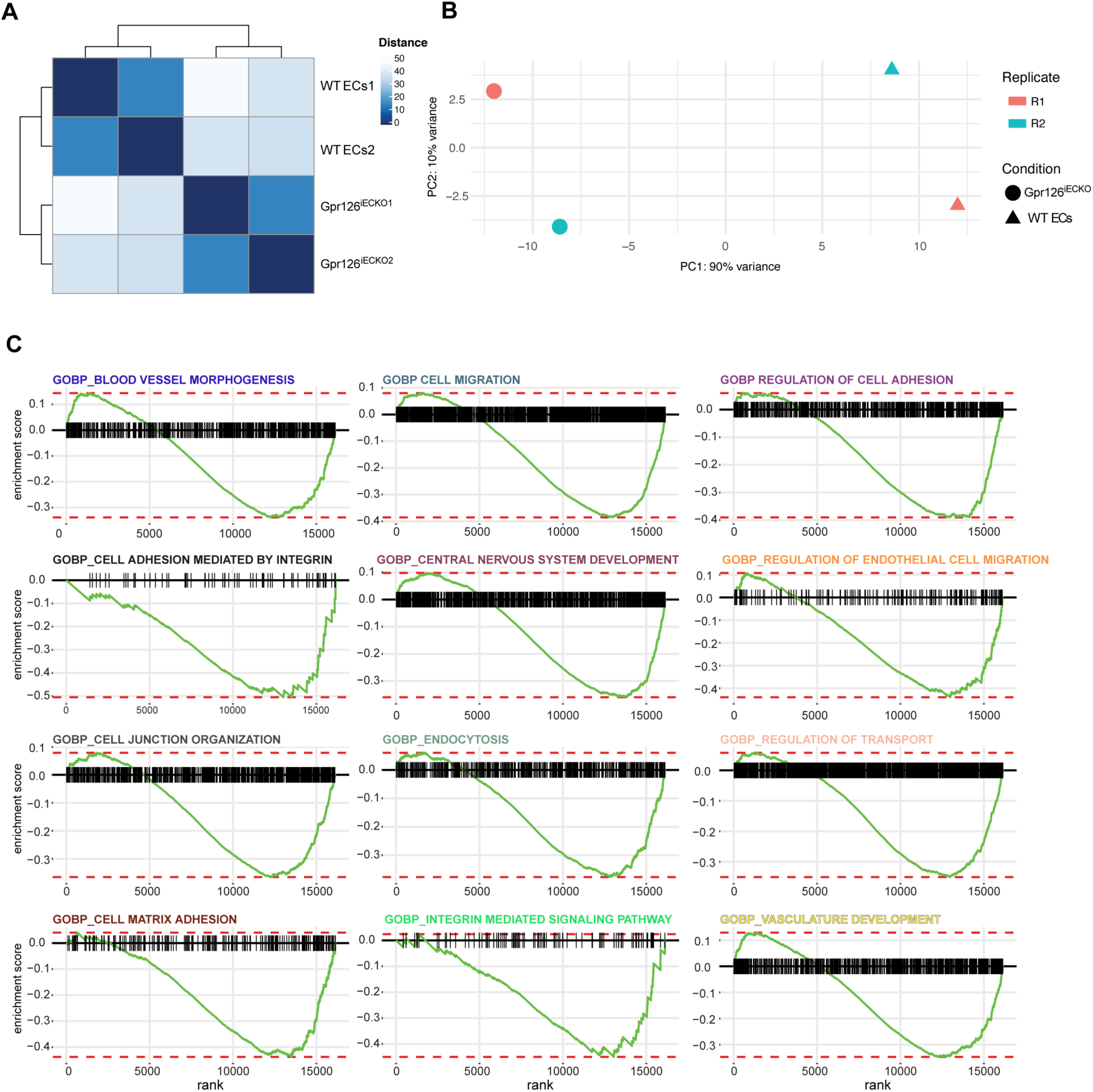
Quality control of RNA-seq data. **A, B, C,** Freshly isolated brain endothelial cells were isolated from WT and Gpr126^iECKO^ mice at P18 and the mRNA was processed for bulk RNA-sequencing. **A,** Distance matrix heatmap showing hierarchical clustering of sample-to-sample distances among the four samples that are the 2 biological replicates of each condition (WT ECs and Gpr126^iECKO^). **B,** Principal component analysis (PCA) plot showing the samples in the 2D plane spanned by their first two principal components. Each symbol represents a biological replicate (R1, R2) for each condition (n=4, 2 WT [circle] and 2 Gpr126^iECKO^ [triangle] mice). **C,** Gene Set Enrichment Analysis (GSEA) plot of selected Gene Ontology Biological Process terms (GOBP). The green curve corresponds to the enrichment score (ES) curve, which is the running sum of the weighted enrichment score obtained from the fGSEA software.

**Supplementary Figure 6.**
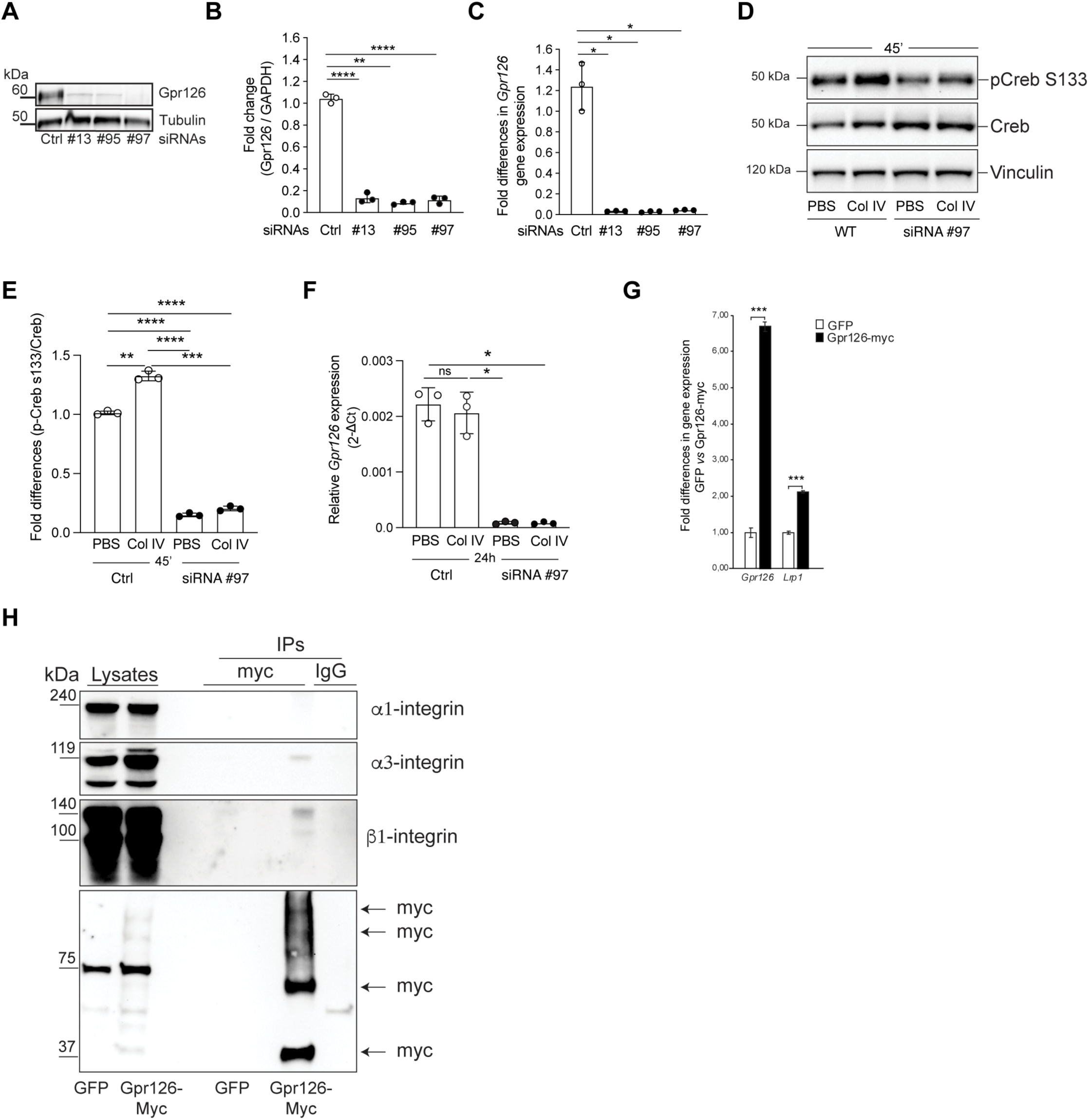
Outline of Gpr126 signaling. **A.** Representative immunoblotting for Gpr126 in cBECs isolated from WT pups at P18 transfected with nontargeting small-interfering RNA (siRNA; Ctrl [control]) or with siRNAs against Gpr126 (no. 13, no. 95, no. 97). Tubulin is shown as loading control. **B,** Gpr126/GAPDH ratio quantified by densitometry scanning and expressed as fold change, as shown in A. Each symbol represents a single experiment (n=12 WT mice for each independent experiment, as means ±SD). **, *P* ≤0.005; ****, *P* ≤0.00005 (Brown-Forsythe and Welch ANOVA, Dunnett’s T3 multiple comparison tests). **C,** Quantification of the relative gene expression of *Gpr126* in cBECs isolated from WT pups at P18, as shown in **A**. Each symbol represents a single experiment (n=12 WT mice for each independent experiment, as means ±SD). *, *P*<0.05; Brown-Forsythe and Welch ANOVA, Dunnett’s T3 multiple comparison tests). **D,** Representative immunoblotting for total Creb and its phosphorylation (phospho-Creb s133) in cBECs isolated from P18 pups, transfected with nontargeting small-interfering RNA (siRNA; Ctrl [control]) or with siRNAs against Gpr126 (no. 97) and treated with Collagen IV (or PBS as vehicle) for 45’. Vinculin is shown as loading control. **E,** p-Creb s133/Creb ratio quantified by densitometry scanning and expressed as fold change, as shown in **D**. Each symbol represents a single experiment (n=12 WT mice for each independent experiment, as means ±SD). **, *P*<0.005; ***, *P*<0.0005, ****, *P*<0.00005 (Brown-Forsythe and Welch ANOVA, Dunnett’s T3 multiple comparison tests). **F,** Quantification of the relative gene expression of *Gpr126* in cBECs isolated from pups at P18, as shown in **D,** except for the treatment with collagen IV for 24h. Each symbol represents a single experiment (n=12 WT mice for each independent experiment, as means ±SD). *, *P*<0.05; ns=not significant (Brown-Forsythe and Welch ANOVA, Dunnett’s T3 multiple comparison tests). **G,** Quantification of fold differences in *Gpr126* an *lrp1* expression in iBECs, following RT-qPCR analysis. (n=3, as means ±SD). ***, *P*<0.0005 (unpaired t-tests with Welch’s correction). **H,** Representative immunoblotting for α1-integrin, α3-integrin, β1-integrin, and myc in iBECs transfected with GFP and Gpr126-myc. The protein extracts (lysates) were immunoprecipitated with an anti-myc antibody [IPs (myc)] or IgG [IPs (IgG)] (only for Gpr126-myc cells), as control. Data are representative of 3 independent experiments.

**Supplementary Figure 7.**
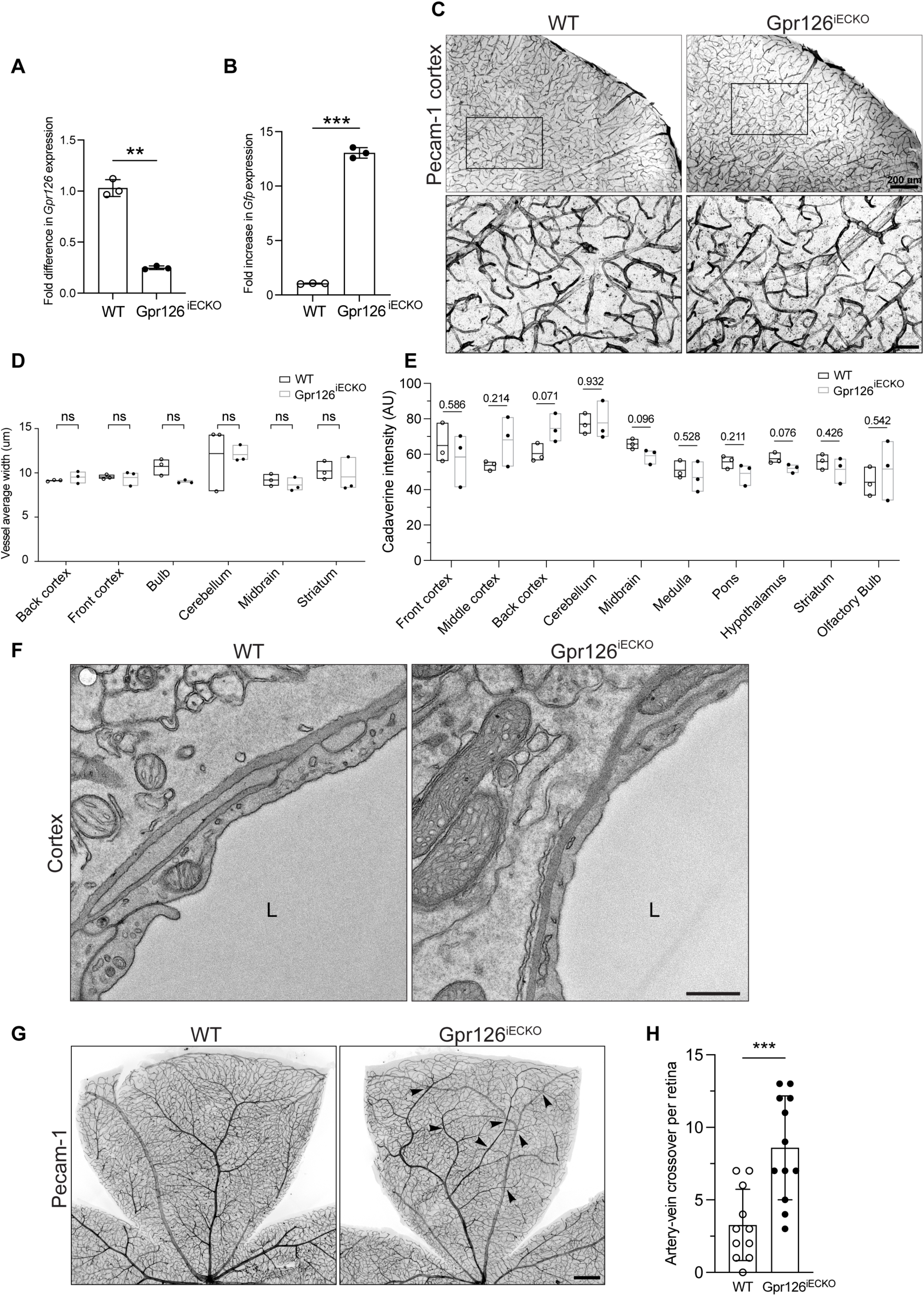
Adult Gpr126^iECKO^ mice have intact BBB although the retinal vasculature shows frequent arteriovenous crossovers. **A, B,** Quantification of the fold differences in *Adgrg6* (**A**) and *Gfp* (**B**) gene expression in fBECs from adult WT and Gpr126^iECKO^. Each symbol represents a mouse. (n=3 WT and 3 Gpr126^iECKO^ mice, as means ±SD). **, *P*<0.005; ***, *P*<0.0005 (unpaired t-tests with Welch’s correction). **D,** Representative confocal images for Pecam-1 in vibratome brain sections (100 μm) from adult WT and Gpr126^iECKO^ mice. Scale bar: 500 μm. Black boxes show cortex regions that are magnified at the bottom. Scale bar: 50 μm. **E,** Quantification of vessels average width in different brain regions of adult WT and Gpr126^iECKO^ mice. Each symbol represents a mouse (n=3 WT and 3 Gpr126^iECKO^ mice, as means ±SD). ns=not significant (unpaired t-tests with Welch’s correction). **E,** Quantification of cadaverine leakage in different brain regions of adult WT and Gpr126^iECKO^ mice. Each symbol represents a mouse (n=3 WT and 3 Gpr126^iECKO^ mice, as means ±SD). *P* > 0.05 (unpaired t-tests with Welch’s correction). **F,** Representative tomography virtual slices of brain blood capillary longitudinal cortex section from adult WT and Gpr126^iECKO^ mice (n=3, WT; n=3, Gpr126^iECKO^). L = lumen. Scale bar: 500 nm. **G,** Representative confocal images for Pecam-1 in the retina of adult WT and Gpr126^iECKO^ mice. Scale bar: 500 μm. Black arrowhead indicates arteriovenous crossovers. Scale bar: 500 μm. **H,** Quantification of the number of arteriovenous crossovers in in adult WT and Gpr126^iECKO^ retinas, as shown in **E**. Each symbol represents a single retina. (n=11 WT and 12 Gpr126^iECKO^ mice, as means ±SD). ***, *P* ≤0.0005 (unpaired t-tests with Welch’s correction).

**Supplementary Figure 8.**
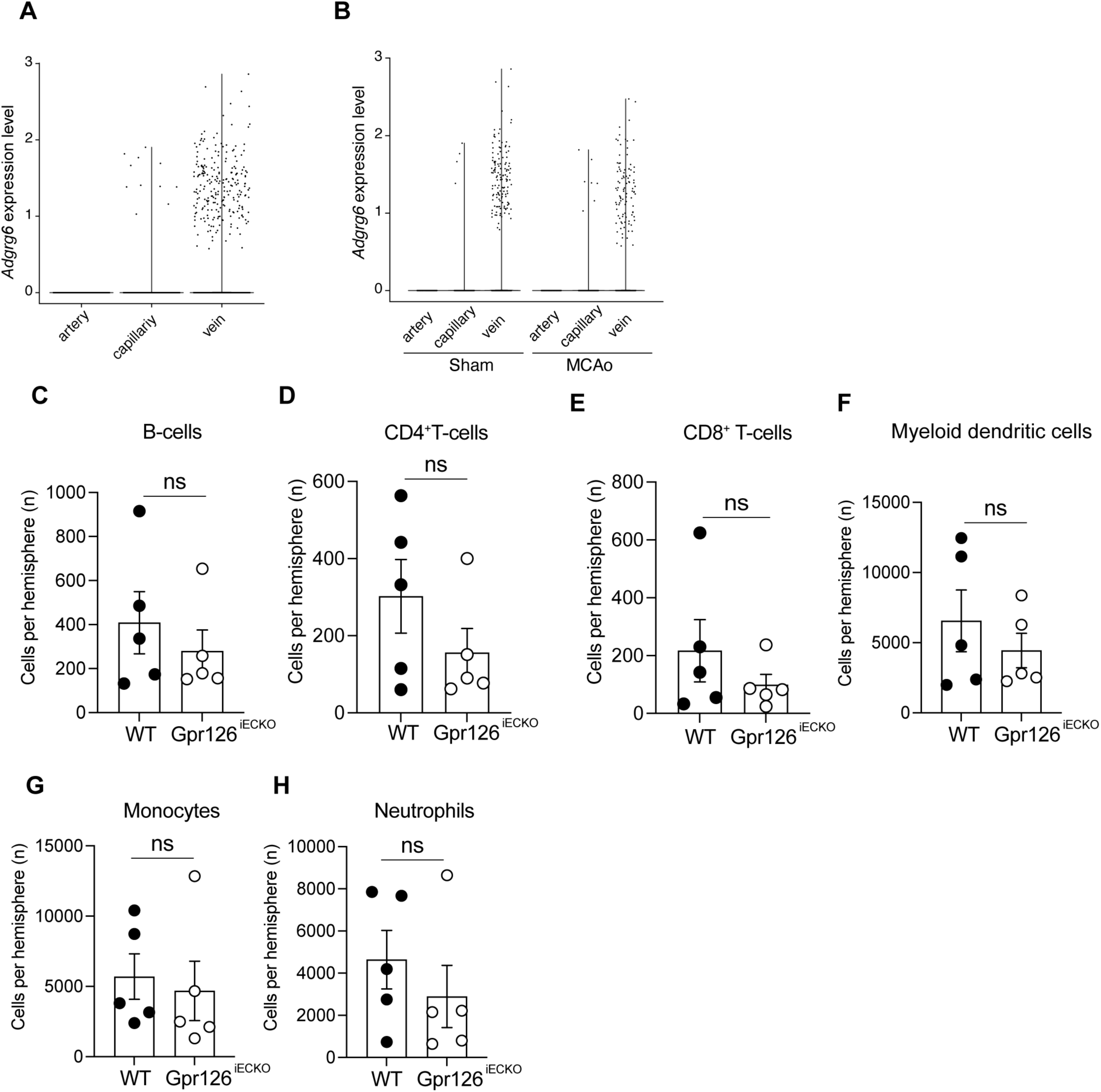
Gpr126 is mainly expressed in vein ECs and no changes in both expression and immune cells’ number were observed after MCAo. **A, B,** Single-cell gene expression profiles of *Adgrg6* (*Gpr126*) in the brain vasculature (artery, capillary and vein) of WT mice. Violin plots displaying the normalized expression values of *Adgrg6* in health (**A**) and after MCAo compared to sham (**B**). Each dot represents a single cell. Data were obtained from Zheng et al. (2022). Cell type annotation was performed according to the expression of the top 50 marker genes reported by Zheng et al. (2022). ECs subtypes were assigned using the top 50 markers genes of brain ECs subtypes of Kalucka et al. (2020). **C, D, E, F, G, H,** Flow cytometric analysis of immune cells in the stroke hemisphere from WT and Gpr126^iECKO^ mice 3 days after MCAo. Quantification of the number of B-cells (**C**) CD4-positive T-cells (**D**) CD8-positive T-cells (**E**) myeloid dendridic cells (**F**), monocytes (**G**) and neutrophils (**H**). Each symbol represents a mouse (n=5 WT and 5 Gpr126^iECKO^ mice for each independent experiment, as means ±SD). ns=not significant (unpaired t-tests with Welch’s correction).

**Supplementary Movie 1. Gpr126 depletion reduced sprouting spheroids in the angiogenesis assay *in vitro***. Time-lapse recording of sprouting spheroids from fBECs of WT and Gpr126^iECKO^ mice at P18, as shown in Figure 4I. Images were acquired by time-lapse widefield microscopy (Thunder Leica) using a 20× objective. Frames were taken every 30 min for 48 h. Arrows show a sprouting EC over time. Arrowheads show a retracting EC over time.

